# Immune Age Acceleration Stratifies Survival in Multiple Myeloma Independent of Chronological Age and Clinical Fitness

**DOI:** 10.64898/2026.11.17.767344

**Authors:** Katherine E. Ferguson, Bridget E. Neary, Sarthak Satpathy, William C. Pilcher, Marina E. Michaud, Denis Ohlstrom, Edward Henderson, Mojtaba Bakhtiari, Steven Foltz, Chaitanya R. Acharya, Immune Atlas Network, Megan M. Weivoda, Ioannis S. Vlachos, David Avigan, Ravi Vij, Samir Parekh, Hearn Jay Cho, Sacha Gnjatic, Li Ding, Mala Shanmugam, Lawrence Boise, George Mulligan, Shaji Kumar, Linda B. Baughn, Ashley Rosko, Sagar Lonial, Ajay Nooka, Manoj Bhasin

## Abstract

Chronological age incompletely explains prognostic heterogeneity in multiple myeloma (MM), particularly among younger patients, whose clinical outcomes vary widely. Existing immune aging clocks are predominantly derived from peripheral blood expression data and thus incompletely reflect the aging of the bone marrow (BM) immune microenvironment in MM. To address this limitation, we developed BM immune aging clocks for ten immune and progenitor cell types trained on single-cell RNA-sequencing data from 1,260 healthy peripheral blood and BM samples and calibrated to BM. We then applied these clocks to BM single-cell profiles from 337 newly diagnosed MM patients enrolled in the CoMMpass study to derive a global immune age acceleration score. In patients younger than 70 at diagnosis, accelerated immune aging was independently associated with shorter overall survival (OS; HR = 2.24, p = 0.003) after adjustment for cytogenetic risk, transplant status, triplet therapy administration, chronological age, sex, performance status, and laboratory deficits, whereas chronological age was not predictive. Subsequent analyses identified CD4+ memory T cell aging as the strongest contributor to overall immune age acceleration and its association with shorter OS. Consistent with this, accelerated immune aging was characterized by terminal effector differentiation of memory T cells, TNF-α/NF-κB-mediated inflammation, and increased lymphocyte-driven TNF, CCL3, and IFN-γ signaling. However, inflammatory signaling alone did not account for survival differences. Longitudinally, changes in monocyte immune age acceleration at first progression were associated with shorter subsequent survival (HR = 3.06, p = 0.005). Together, these findings identify BM immune age acceleration as a prognostic measure of biological fitness in MM, distinct from chronological age and clinical frailty.

## INTRODUCTION

Multiple myeloma (MM) is a cancer of plasma cells that develops and progresses within the bone marrow (BM).^1^ MM is predominantly a disease of older adults, with a median age of 69 years at diagnosis, though its incidence in patients under age 40 has nearly doubled over the past three decades.^2^ Chronological age is a well-established independent prognostic factor in MM, with patients under the age of 60 exhibiting nearly twice the 5-year survival rate of those over 70 years old.^3, 4^ Furthermore, even a single-year increase in age has been associated with a 4% to 8% increase in the hazard of death.^5^ However, younger patients still exhibit significant heterogeneity in their disease progression and outcomes.^6^

Chronological age alone fails to capture the molecular and physiological heterogeneity driving these differences in outcomes, particularly in younger patients receiving autologous stem cell transplantation (ASCT), for whom age fails to stratify risk.^7^ To address this, clinicians rely on measures of performance status, such as Eastern Cooperative Oncology Group (ECOG) performance status, Karnofsky Performance Status (KPS) or frailty metrics, to stratify risk beyond chronological age alone.^8, 9^ Yet these measures are often subjective, rely on cumulative deficits and may have limited discriminatory value in younger transplant-eligible patients, who may have substantial underlying differences in biological fitness but similar functional status. Therefore, measures of biological fitness that complement age, performance status, frailty indices, and established clinical risk factors are needed to more accurately predict disease progression and treatment response in MM.

Immune aging, characterized by immunosenescence and inflammaging, is a prominent feature of biological aging. In MM, age-related decline in adaptive immunity and chronic low-grade inflammation have been associated with disease progression and reduced therapeutic efficacy.^10,11^ T cell dysfunction, increased inflammatory cytokine production, and shifts in immune cell populations can all facilitate immune evasion by malignant plasma cells. In addition, these changes can impair immunotherapies that rely on host immune fitness for efficacy, such as CAR-T cells, bispecific antibodies, and checkpoint inhibitors.^11, 12^

The recent development of immune aging clocks using transcriptomic and epigenetic data has advanced researchers’ ability to quantify biological age.^13, 14^ These immune age clocks, however, are trained predominantly on peripheral blood (PB) tissue. In hematological malignancies, particularly MM, the fitness of the local BM immune microenvironment is critical.^15^ Moreover, the circulating immune system is highly dynamic, with leukocyte composition and transcriptional states fluctuating with circadian rhythm, acute stress, and pharmacological intervention, such that a single blood draw captures a transient snapshot rather than the stable, tissue-resident immune landscape.^16, 17^ Therefore, measuring biological age using blood-based immune aging clocks does not capture the full extent of immune dysfunction most relevant to disease biology and patient outcomes in MM.

To address this gap, we developed bone marrow-based aging clocks for ten major immune and progenitor cell types, trained on single-cell RNA-sequencing (scRNA-seq) data from healthy PB and BM donors, and combined them into a global immune age score. We applied these clocks to our comprehensive MM immune atlas, including 337 newly diagnosed CoMMpass patients who have well-characterized baseline data and long-term clinical outcomes as well as paired follow-up samples. We used global immune age acceleration to assess patient risk at diagnosis and cell type-specific clocks to identify the transcriptional programs associated with immune aging in each compartment, capturing cell type-specific changes that may not be reflected in the global score, including those following treatment. We then systematically evaluate how accelerated immune aging shapes the tumor microenvironment and influences patient outcomes.

## METHODS

### 1. Data Acquisition and Cohort Assembly

To construct the comprehensive single-cell RNA sequencing atlas, healthy peripheral blood and bone marrow samples were compiled from 12 public datasets, including 8 from the DISCO database (E-MTAB-8884, E-MTAB-9139, GSE120221, GSE130430, GSE135194, GSE159929, GSE175604, GSE188222), the Human Cell Atlas, GSE299043, the OneK1K cohort (GSE196830), and the Sage blood atlas (syn49637038). Donors younger than 18 years were excluded. Peripheral blood donors were included to increase the number of training donors and to cover ages that were sparse or tied to a single dataset among bone marrow donors. Models were weighted toward and calibrated to bone marrow (see Model Training and Bone Marrow Calibration). Supplemental Table S1 lists each dataset with accession ID number, tissue, number of donors, age range, and chemistry.

### 2. Quality Control, Integration, and Annotation

Datasets were processed using Scanpy (version 1.11.5). Within each dataset, cells with fewer than 100 detected genes and genes detected in fewer than three cells were removed. In the merged atlas, cells were additionally filtered by >200 detected genes, ≥1,000 and ≤80,000 UMIs, <15% mitochondrial and <15% hemoglobin reads. Doublets were identified per sample with scDblFinder (1.22.0) in all datasets except Onek1k, which had been quality-controlled by the original study and used as distributed. Clusters enriched for predicted doublets and with mixed-lineage signatures were removed during annotation. Counts were normalized to 10,000 per cell and log1p-transformed.

For annotation, Harmonypy 0.2.0 was run on the first 50 principal components of 2,000 highly variable genes, with the batch key set to the sequencing batch (Sage atlas batch, DISCO accession, HCA sample, or GSE299043 tissue). The Harmony embedding was used for the neighbor graph, clustering, and UMAP. All clocks were trained and applied to log-normalized expression. Cells were assigned to 10 populations using canonical marker expression and original dataset annotations: CD4+ and CD8+ naïve and memory T cells, natural killer (NK) cells, memory and naïve/transitional B cells, monocytes, dendritic cells (DCs), and hematopoietic stem cells (HSCs).

### 3. Donor-Level Train and Test Split

For each of the 10 cell types, donors were split into training (70%) and test (30%) sets using scikit-learn train_test_split (random seed set to 42), so that no donor contributed to both sets within a cell type model. Stratification used age bins (<30 years, 30-65 years, and >65 years), tissue (blood or bone marrow), and sex, with unknown sex treated as a separate category. If any stratum had too few donors for a stratified split, sex, then tissue, and finally age bin was dropped from stratification. If no stratification succeeded, donors were split at random. Strata containing fewer than three donors were assigned entirely to the training set. The stratification used for each cell type is listed in Supplemental Table S2.

Because splits were cell type-specific, a donor could be assigned to the training set of one lineage and the test set of another. Global immune age was therefore evaluated in two ways. For Figure 1D-1E, bone marrow donors with at least 75% of their lineages in training were designated global training donors (n = 51), and the remaining donors were designated global test donors (n = 35). As a leakage-free alternative, each bone marrow donor’s global score was also calculated using only lineages that were held out from training (Supplemental Table S3E).

**Figure 1.**
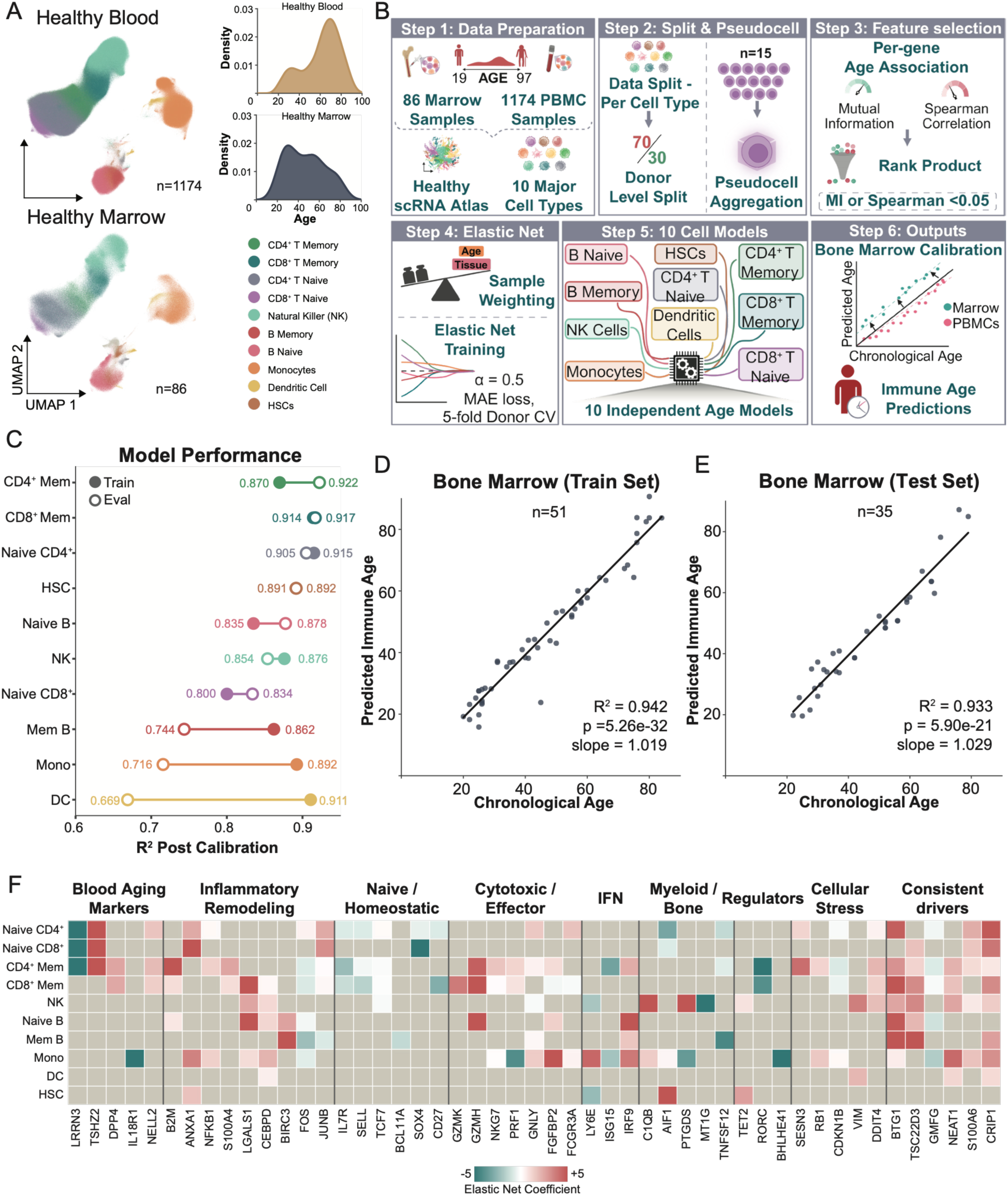
Construction and calibration of a bone marrow-calibrated immune aging clock from healthy single-cell transcriptomes. **(A)** UMAPs and density plots of the reference healthy immune atlas built from 12 publicly available datasets. The top UMAP and density plot show healthy peripheral blood (n = 1174), while the bottom shows healthy bone marrow (n = 86). Cells in UMAPs are colored according to lineage. The distribution of chronological age across donors is plotted, with age on the x-axis and donor density on the y-axis. **(B)** Overview of immune age prediction model design. **(C)** Dumbbell plot showing training and held-out evaluation performance (coefficient of determination, R^2^) of calibrated lineage-specific clocks in bone marrow donors. Training R^2^ values are indicated by filled circles and held-out test R^2^ values are open circles. **(D, E)** Predicted global immune age versus chronological age across healthy bone marrow donors in the training cohort (**D**, n = 51) and held-out test cohort (**E**, n = 35). Donors were assigned to the global training cohort if at least 75% of their cell lineages were included in the training set; all remaining donors were assigned to the global test cohort. Each point represents one donor. The solid line represents the linear regression fit. Model performance is summarized by the regression R^2^, slope, and p-value. **(F)** Tile plot with selected genes (columns) contributing to elastic net models across lineage models (rows), grouped by category. Red represents positive elastic net coefficients (higher expression associated with older predicted age) with max color at +5, and teal represents negative elastic net coefficients with max color at –5. Gray tiles indicate genes not used in that lineage’s model.

### 4. *Pseudocell Creation* and Feature Selection Pre-filtering

To reduce single-cell sparsity while preserving donor variance, each donor’s cells were randomly pooled into groups of approximately equal size (15-29 cells; donors with fewer than 15 cells formed a single group), and log-normalized expression was averaged within each group to form pseudocells. Genes with zero variance across training pseudocells were removed. Feature selection was optimized independently for each lineage utilizing mutual information and Spearman correlation alongside a 100-iteration bootstrap incorporating 80% donor subsampling. For each metric, genes were ranked within each iteration, and ranks were combined across iterations by rank product using geometric mean rank. Genes ranking in the top 5% by rank product for either metric were retained. Depending on lineage, feature selection was either applied globally across the training set or stratified to capture features specific to age groups, including donors under 35 years of age or donors over 65 years of age (Supplemental Table S2).

### 5. Model Training

The core framework for pseudobulking and rank-product feature selection was adapted from the foundational scImmuAging pipeline.^14^ To accurately model the localized bone marrow microenvironment and ensure stringent predictive generalizability, we implemented several critical algorithm modifications. First, prior to finalizing the model architecture, we evaluated alternative linear and non-linear algorithms, specifically Lasso regression and regression splines. An elastic net regression framework, implemented in R’s glmnet (version 4.1.10) (mixing parameter α = 0.5), was used for all final models with λ set to minimize the mean absolute error (MAE) in cross-validation. Second, to prevent intra-donor data leakage during hyperparameter tuning, we explicitly grouped cross-validation folds (n = 5) by donor identity to ensure no donor spanned both training and validation strata. Finally, to handle the unique class imbalances of our cross-tissue atlas, we replaced simple age-based weighting with a multiplicative sample weight ( *w_final_* ) on each pseudocell prior to training. The base weight inversely penalizes overrepresented donors and overrepresented age groups (5-year age intervals). This base weight was multiplied by *w*_>65_ for donors over the age of 65 years and by *w_BM_* for bone marrow, and weights were normalized to the sum of training pseudocells in each lineage. For each cell type, the feature selection strategy, *w_BM_* , and *w*_>65_ were chosen on MAE and R^2^ in bone marrow training (Supplemental Table S2). A donor’s predicted age was the mean prediction across its pseudocells.

### 6. Bone Marrow Calibration

Predicted ages were calibrated to healthy bone marrow. For each cell type, the prediction residual (predicted − chronological age) was regressed on chronological age in bone marrow training donors, giving an intercept (α) and slope (β). Age acceleration and adjusted age were then defined as:

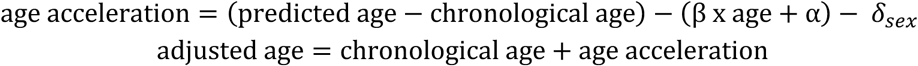

The sex offset δ was estimated in held-out bone marrow donors using a linear mixed model of age-corrected residuals on sex, cell type and age with a random intercept per donor (observations with |z| ≥ 2.5 within cell type excluded). A sex × cell type interaction was not supported (likelihood-ratio test, p = 0.17), so a uniform offset was applied (δ = +1.32 for males and -1.32 for females; δ = 0 for samples of unknown sex). Before correction, male acceleration exceeded female acceleration by 2.64 years.

We assessed calibration using the slope and R^2^ of adjusted predicted age versus chronological age. Because all held-out bone marrow donors originally contributed to the delta parameter, we reported held-out bone marrow and training marrow uncalibrated performance separately (Supplemental Table S3A). Cox model estimates remained unchanged without sex offset (δ). Throughout, age acceleration refers to the calibrated residual.

### 7. Clock Performance

Performance was reported on uncalibrated predictions in bone marrow training donors, bone marrow held-out donors, and peripheral blood held-out donors. Metrics included the squared Pearson correlation (R^2^), MAE, and slope of prediction on chronological age, with 95% CIs from 2,000 donor bootstrap resamples. Within-dataset performance was also assessed by R^2^ within each dataset with ≥8 donors. Dataset effects were also assessed with a linear mixed model of uncalibrated predicted age on chronological age.

### 8. *Global* Immune Age Scoring

Global age acceleration was computed as the weighted mean of the calibrated cell type accelerations, taken from cell types present in each sample:

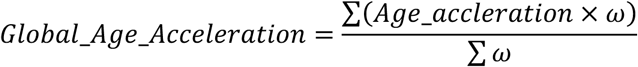

Global immune age was defined as chronological age plus global age acceleration. Lineage weights (ω) were defined as ω= R^2^/(var_accel+25). R² is the squared Pearson correlation between uncalibrated predicted age and chronological age in bone marrow training donors. Var_accel is the variance, in the same donors, of the residual from a regression of uncalibrated predicted age on chronological age. The constant 25 years² (SD of 5 years) limits the weight given to lineages with small residual variance.

### 9. Immune Atlas Cohort

The scRNA-seq cohort was described by Pilcher et al^18^. Briefly, 337 CD138-negative MM bone marrow mononuclear cell (BMMC) samples were collected from patients enrolled in the MMRF CoMMpass study (NCT01454297). Patients enrolled in the study were regularly monitored via quarterly check-ins for up to eight years following initial diagnosis. Baseline samples were acquired prior to the first-line therapy. MMRF CoMMpass data can be accessed through the MMRF VLAB upon approval (https://mmrfvirtuallab.org).

### 10. Application of Clocks to Immune Atlas Cohort

Single-cell annotations identified in the Pilcher et al.^18^ immune atlas were mapped to the 10 clock lineages (Supplemental Table S5). Plasma cells, erythroid cells, granulocytes, fibroblasts, and megakaryocytes were excluded. Lineage clocks were applied to each sample and calibrated as above, using age at diagnosis and recorded sex. Patients were analyzed in two age groups, <70 and ≥70 years, reflecting a threshold commonly used for ASCT eligibility in clinical trials.^19^ Although ASCT eligibility is now based on fitness rather than age alone, this cutoff divided the cohort into groups that differed in first-line treatment (ASCT 59% versus 32%; triplet induction 88% versus 65%) and in whether chronological age stratified survival. Within each age group, patients were divided into quartiles of global age acceleration (Q1, slowest; Q4, fastest). Lineage-specific quartiles were defined within each cell type and age group. At diagnosis, global age acceleration was the primary measure for patient stratification, and cell type and subtype analyses were used to evaluate what drives global age acceleration. In follow-up samples, where cell type accelerations were less correlated with one another, cell type clocks were also analyzed individually. Patients in the highest and lowest quartiles of global age acceleration are referred to as fast and slow agers, and patients in the highest and lowest quartiles of a cell type-specific acceleration are referred to as having the most and least accelerated cells of that type.

### 11. Survival Analysis

PFS and OS were analyzed with Cox proportional hazards models. Primary models included global age acceleration (per 10 years), age at diagnosis, sex, IMWG high risk, unknown IMWG risk, first-line ASCT and first-line triplet therapy, stratified by collection site. Models were fit separately for patients <70 and ≥ 70 years. Each model used all patients with complete data for model variables. Proportional hazards were assessed with Schoenfeld residual tests (Supplemental Tables S8 and S19). As sensitivity analyses, models were refitted with bone marrow plasma cell percentage (per 10%) and scRNA-seq monocyte (per 10%) and DC (per 1%) proportions of immune cells added. Kaplan-Meier curves compared Q1 and Q4 within each age group with log-rank tests. Hazard ratios for Q4 versus Q1 were adjusted for ASCT. Hazard ratios were also expressed per standard deviation of global age acceleration. To test whether any single lineage drove the global age acceleration association, global age was recomputed leaving one lineage out at a time and refit on primary models. Cell type-specific accelerations (per 10 years) were modeled one at a time with the same covariates and site stratification, with Benjamini-Hochberg correction across the five T and NK lineages, and separately, across the five B cell, myeloid and progenitor lineages. Effect modification by age group was tested with an age acceleration x age group (≥70 versus <70) interaction term in models including both age groups. To assess survival outcomes with chronological age, Kaplan-Meier curves by decade (<50, 50-59, 60-69, 70-79 and ≥80 at age of diagnosis) were compared within each age group with multivariate log-rank tests. Kaplan-Meier curves were truncated when fewer than three patients were at risk, and numbers at risk are reported in Supplemental Table S9.

### 12. Cell Composition and Clinical Associations

Lineage proportions were calculated as the fraction of all immune cells. Pairwise correlations of lineage age acceleration were computed with Pearson correlation in patients with all 10 lineages, with Benjamini-Hochberg (BH) correction. Plasma cell percentage was compared between Q1 and Q4 with a Mann-Whitney U test and correlated with global age acceleration using Spearman correlation. Lineage proportions were also correlated with global age acceleration using Spearman correlation.

### 13. Gene Expression, Pathway, and Transcription Factor Analysis

We used two complementary analyses for differential expression. We performed a bystander analysis of genes associated with predicted age (Figures 3A and 4A) and a differential expression analysis comparing patients with the highest and lowest age acceleration for each lineage clock (Figures 3B and 3I; Supplemental Figures 3G and 4D).

#### 13.1. Bystander analysis

For each lineage, log-normalized gene expression was averaged across cells within each donor or patient. The MM cohort included patients younger than 70 years with a clock prediction for that cell type (at least 15 cells), and the healthy reference cohort included all healthy bone marrow donors in the atlas, including training donors (Supplemental Tables S2 and S5). Genes used by that cell type’s clock were removed so that only bystander genes were tested. Remaining genes were ranked by partial Spearman correlation with predicted age, adjusted for collection site, sex, and log median genes per cell in MM and for log median genes per cell in healthy donors. Correlations were Fisher z-transformed for gene ranking.

#### 13.2. Most Versus Least Accelerated Analysis

For each cell type, raw counts were pseudobulked per patient (aggregate) in patients younger than 70 years, and patients in the highest (Q4) and lowest (Q1) quartiles of that cell type’s age acceleration were compared. Pseudobulk samples with fewer than 15 cells or 1,000 counts were removed. Differential expression was tested with pyDESeq2 using the design ∼ site + sex + log median genes per cell + quartile. Wald statistics were used as gene-level statistics. Clock genes were retained in this analysis. Gene-level, pathway, and transcription factor results are in Supplemental Table S11.

#### 13.3. Pathway and Transcription Factor Analysis

For both DEG analyses, pathway and transcription factor activity were inferred from the gene-level statistics with the univariate linear model (ULM) from decoupler for MSigDB Hallmark, MSigDB Reactome, and CollecTRI. Reactome pathways related to infection or neurological pathways and pathways defined by a specific mediator (containing BY, MEDIATED, IN, DURING, or DEPENDENT) were removed, and only Reactome pathways with 25-200 genes were retained. P values were Benjamini-Hochberg corrected for each lineage, cohort, and gene selection set. Significance was defined as adjusted p<0.05, and adjusted p<0.1 was shown *as a trend in dot plots*.

### 14. NF-kB and Clock Comparison

An NF-κB activity score was computed per patient from Hallmark TNF-α signaling via NF-κB, excluding clock genes on pseudobulked NK, CD4^+^ and CD8^+^ naïve and memory T cells. Nested Cox models with the same covariates as the primary survival models compared NF-κB alone, the clock alone, and both, using likelihood-ratio tests. Aggregate NK/T age acceleration was defined as the weighted mean of the five T and NK clock accelerations using their global weights. The NF-κB score was modeled per standard deviation.

### 15. Shift-Share Decomposition

For compartments with more than one subtype, each patient’s age acceleration was decomposed into a composition component (differences in subtype proportions), a transcriptional component (differences in subtype-level acceleration), and their interaction based on shift-share decomposition.^20^ Decomposition was restricted to subtypes with accelerations in ≥80% of patients and ≥10 patients overall. Subtype-level acceleration, *a_is_*, was computed by applying a lineage-specific clock for each subtype, *s*, to mean log-normalized pseudobulk expression. Subtype proportions, *p_is_*, were renormalized to the included subtypes with ≥80%, and missing subtype accelerations were imputed using the cohort mean *ā_s_*. The composition-weighted overall acceleration, *A* = ∑*_s_ p_is_a_is_*, was decomposed around the cohort means *ā_s_* and *p̄_s_* using the equation below:

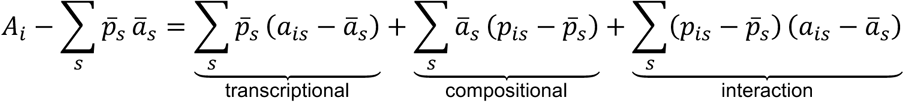

The three components were then z-scored and entered jointly into a Cox model for OS, adjusted for age, sex, ASCT, IMWG high and unknown risk, and first-line triplet therapy, stratified by site in patients <70 years old. Each component’s relative contribution to the mortality hazard was determined by its share of the variance of the linear predictor, 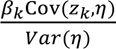.

### 16. Subtype Abundance, Trajectories, and Subtype-Level Clocks

Subtype annotations were taken from Pilcher et al.^18^ Subtype proportions within each compartment were compared between patients with the most and least accelerated cells for that compartment’s clock using a Mann-Whitney U test with Benjamini-Hochberg correction (Figure 3F-G; Supplemental Figures 3F and 4C).

For memory T cells, differential abundance was displayed on a trajectory constructed with Slingshot (version 2.10.0) on the first 25 Harmony-corrected principal components computed across all CD4^+^ and CD8^+^ T cells, with naïve CD4^+^ and naïve CD8^+^ clusters as start clusters, respectively. Trajectory connectivity from Slingshot was converted to a tree graph with igraph (version 2.0.3). Node size shows the number of cells in each cluster, and node color shows the differential abundance as log2 fold change in mean proportion Q4 versus Q1.

To test subtype-level clocks (Figure 3H), the parent cell type clock was applied to mean log-normalized pseudobulk profiles of each subtype in patients under 70 and with at least 10 cells of that subtype, and predictions were calibrated with the parent cell type’s bone marrow calibration. Patients were divided into quartiles of subtype-level accelerations, and the most and least accelerated groups of that subtype were compared with Kaplan-Meier curves and log-rank tests in subtypes with at least 40 patients. All subtypes tested are in Supplemental Tables S2 and S5.

### 17. Follow-Up Samples

Paired baseline and follow-up bone marrow samples were available for 72 patients younger than 70 years at baseline (85 pairs). Follow-up samples were grouped as collected after induction chemotherapy (IC), after induction followed by ASCT (IC + ASCT), or at first progression based on patient visit tables collected by CoMMpass. Follow-up samples were annotated and scored with the same lineage clocks and calibration as baseline samples using age at the follow-up draw. The change in age acceleration (Δ) was calculated as follow-up minus baseline acceleration for each lineage and for global age acceleration and was tested against zero within each group with Wilcoxon signed-rank tests. Comparisons among cell types were summarized by Pearson correlations of lineage accelerations and by the within-patient standard deviations across lineages, using a two-sided Wilcoxon signed-rank test. Sample counts, Δ summaries, residualization fits and survival models for the follow-up cohort are in Supplemental Table S14.

### 18. Residualized Change in Monocyte Age Acceleration

Since Δ monocyte acceleration at first progression depended on baseline acceleration, Δ was residualized on baseline monocyte acceleration and time between draws with linear regression. The residual was used to rank patients. OS was measured from the progression draw. Cox models included residualized Δ (per 10 years), age, IMWG high risk and unknown risk, sex, and first-line ASCT, stratified by site, with a ridge penalty of 0.1. First-line triplet therapy was omitted because 42 out of 44 patients received it. Kaplan-Meier curves compared patients above and below the median residualized Δ with a log-rank test. Genes associated with residualized Δ were estimated in monocyte pseudobulk profiles at first progression with pyDESeq2 using the design ∼ site + sex + log median genes per cell + Δ mono residual per 10 years. Genes were retained if they had at least 10 counts in at least 25% of samples. Wald statistics were used as gene-level statistics. Pathway and transcription factor activity were inferred with ULM as above. Baseline-to-progression changes were estimated in paired monocyte pseudobulk profiles with a patient term separately for patients above and below the median residualized Δ. The difference between groups was tested with a timepoint x group interaction, adjusted for the interaction of timepoint with site, IMWG risk, first-line ASCT, time between draws, and sequencing depth. Gene filtering and ULM pathway analysis were as above.

### 19. Performance Status and Laboratory Deficit Score

ECOG performance status at diagnosis was grouped as 0, 1, or ≥2. The laboratory deficit score (LDS-5) was built from five baseline laboratory measures: blood urea nitrogen (BUN), glucose, platelet count, white blood cell count (WBC), and body mass index (BMI). Each measure was scored as one deficit when outside of the following thresholds: BUN > 20 mg/dL, glucose <70 or >140 mg/dL, platelets <150 or > 450 x10^9^/L, WBC <4.5 or >11 x10^9^/L, and BMI <18.5 or ≥30 kg/m^2^. Candidate laboratory values were excluded if they contributed to established disease burden measures such as ISS, R-ISS, or CRAB criteria: albumin, creatinine, corrected calcium, LDH, hemoglobin, and β2-microglobulin. Lab values were also excluded if they had more than 40% missing (C-reactive protein) or were redundant with another laboratory value (absolute neutrophil count; Supplemental Table S17). LDS-5 was the number of deficits (0-5), prorated to five items for patients with four of five measures available, and was grouped as 0,1, or ≥2 for Kaplan-Meier analysis.

For ECOG and LDS-5, survival across categories was tested with a univariate Cox likelihood-ratio trend test treating category as ordinal, and each category was compared with the lowest category using log-rank tests with Holm correction.

Associations between immune age acceleration, ECOG, LDS-5, disease burden measures (ISS stage, β2-microglobulin, creatinine, albumin and LDH), and quality-of-life (QOL) measures (using the European Organization for Research and Treatment of Cancer [EORTC] QLQ-C30 domains global health status, physical functioning, fatigue and pain) were estimated with partial Spearman correlations with Benjamini-Hochberg corrections adjusted for chronological age, separately in each age group.

### 20. Joint Models of Immune Age Acceleration, ECOG, and LDS-5

Immune age acceleration (per 10 years), ECOG (per category), and LDS-5 (per standard deviation) were entered together in Cox models adjusted for age, sex, IMWG risk, first-line ASCT and first-line triplet therapy, stratified by collection site. These models were restricted to patients with complete data for all variables, and each measure was also modeled alone in the same patients. Age-group interactions were tested for each measure. Sensitivity analyses added ISS stage, used missing indicator terms for ECOG and LDS-5 to retain all patients, and compared immune age acceleration between included and excluded patients with a Mann-Whitney U test.

### 21. Cell Communication

Cell communication was inferred with LIANA (version 1.10) using its consensus ligand-receptor resources in two complementary ways:

#### 21.1. Per-condition analysis

LIANA rank_aggregate was run separately on cells from fast (Q4) and slow (Q1) agers. For each sender-receiver cell type pair and ligand-receptor pair, we evaluated the mean ligand-receptor expression (lr_means, a measure of interaction strength) and a CellPhoneDB permutation p-value for cell type-specificity (p<0.05 threshold). An interaction is absent from a quartile’s output and considered not detected there because LIANA removes pairs whose ligand or receptor is expressed in fewer than 10% of cells of the sender or receiver.

#### 21.2. Differential-expression-based analysis

Pseudobulk profiles (patient × cell type) were tested for fast (Q4) versus slow (Q1) agers with PyDESeq2 (design ∼ site + sex + log median genes per cell + global quartile). Differentially expressed genes were mapped to ligand-receptor interactions to identify deregulated intercellular communication pathways. An interaction was kept when the ligand and receptor were both significant (adj. pdj < 0.05) and changed in the same direction, or when one partner was significant and the other was not. Interactions were then verified against the per-condition runs: the interaction had to be specific (CellPhoneDB p < 0.05) in the quartile where DE predicted it to be higher and have higher lr_means in that quartile. Downstream responses in receiver cells were estimated per cell with the univariate linear model (ULM) of decoupler (version 2.1.2) on log-normalized expression. Three prior-knowledge resources were used: PROGENy pathway footprints (top 500 genes per pathway), CollecTRI transcription factor regulons, and the MSigDB Hallmark gene sets. To separate type II from type I interferon responses, we derived two subsets of Hallmark genes: IFN-γ-only (genes in HALLMARK_INTERFERON_GAMMA_RESPONSE and not in HALLMARK_INTERFERON_ALPHA_RESPONSE) and IFN-α-only (the reverse). Sources with fewer than five target genes in the data were not scored. Up to 300 randomly sampled cells per patient × cell type (fixed seed) were scored, and per-cell scores were averaged per patient × cell type (≥ 10 cells).

### 22. Statistical Analysis and Software

All tests were two-sided. Unless otherwise stated, p<0.05 was considered significant, and multiple comparisons were corrected with the Benjamini-Hochberg method. Analyses were performed with R (version 4.5.1), glmnet (version 4.1.10), infotheo (version 1.2.0.1), scDblFinder (version 1.22.0), slingshot (version 2.10.0), and igraph (version 2.0.3) packages. Python (version 3.10.19) with Scanpy (version 1.11.5), harmonypy (version 0.2.0), scikit-learn (version 1.7.2), decoupler (version 2.1.2), PyDESeq (version 0.5.2), lifelines (version 0.30.0), LIANA (version 1.10), statsmodels (version 0.14.6), and SciPy (version 1.15.3) were also used.

## RESULTS

### Development and validation of bone marrow immune aging clocks

To develop immune aging clocks applicable to the MM bone marrow microenvironment (BMME), we assembled a comprehensive healthy donor scRNA-seq reference atlas comprising 86 healthy BM and 1,174 PB samples from individuals aged 19 to 97 years (**Figure 1A**). Given the limited availability of healthy BM samples, we incorporated PB samples to expand the training cohort, while assigning greater weight to BM samples in some cell types during model development. For each of the ten major immune and progenitor populations in the BMME, we trained an elastic net regression model to predict chronological age from cell type-specific transcriptional profiles (*see Methods*). This produced individual immune aging clocks for CD4^+^ memory T cells, CD8^+^ memory T cells, naïve CD4^+^ T cells, naïve CD8^+^ T cells, natural killer (NK) cells, memory B cells, naïve B cells, monocytes, dendritic cells (DCs) and hematopoietic stem and progenitor cells (HSCs), whose scores were subsequently combined into a global immune aging score (**Figure 1B**).

We first evaluated the ten immune aging clocks in a test set of held-out healthy donors (**Supplemental Figure 1A**). Across all held-out donors before calibration, predicted ages were positively correlated with chronological age in all clocks (r = 0.50-0.77; MAE = 8.6-12.6 years). However, the slopes of predicted chronological age ranged from 0.297 to 0.572, indicating systematic overestimation of age in younger donors and underestimation in older donors. These associations were also observed in individual datasets, indicating that clock performance was not driven by differences between datasets (**Supplemental Table S3C**). Thus, while the clocks captured age-associated transcriptional changes in the test set, their raw predictions understated age differences between donors.

Given the predominance of PB samples in the training cohort, we next evaluated whether the clocks retained their predictive performance in the BM samples of our test set. Despite the smaller number of BM donors, correlations between predicted and chronological age were higher in BM than PB for seven out of the ten cell types (**Supplemental Figures 1B-C**). In held-out BM samples, positive correlations were significant for nine cell type-specific clocks (r = 0.53-0.93; MAE = 4.82-11.1 years). The DC clock did not show a significant association with age, which may be due to fewer BM donors available for evaluation (**Supplemental Table S3A**). In PB, all ten clocks were positively correlated with chronological age (r = 0.54-0.75), although prediction errors were larger (MAE = 9.0-12.4 years). Together, these results demonstrated that the cell type-specific clocks captured age-related variation in BM despite being trained predominantly on PB, but calibration was needed to account for the systematic underestimations in older donors and overestimations in younger donors.

To remove this compression toward the mean, we calibrated each clock using healthy BM samples so that its output could be interpreted as “age acceleration”, defined as the difference between a donor’s predicted age and their chronological age. A positive value, therefore, indicates a donor’s immune cells appear older than expected, whereas a negative value indicates a younger-than-expected immune cell profile. In the BM test set, the regression slopes of calibrated predictions on chronological age ranged from 0.86 to 1.14 for nine cell types, with a lower slope for DCs (0.65; **Supplemental Figure 1D**), and R^2^ ranged from 0.669 to 0.922 (**Figure 1C**), reflecting how widely calibrated predictions varied around chronological age in each cell type. This calibration makes age acceleration directly comparable between patients of different chronological ages.

To summarize immune aging across cell types into a single patient-level measure, we combined the calibrated cell type clocks into a global immune age score. Global immune age, defined as chronological age plus weighted age acceleration score, demonstrated excellent performance on training (R^2^ = 0.942; MAE = 3.42 years) and test sets (R^2^ = 0.933, MAE = 3.77 years; **Figure 1D, 1E**). Because training and test groups differed across cell types, some donors contributed to the training of certain clocks while being held out from others. To account for this overlap, we further evaluated the global immune age score using only cell type clocks that had excluded each donor during training (n = 58). In this analysis, uncalibrated predictions remained correlated with chronological age (R^2^ = 0.701; MAE = 7.89 years) and calibrated global immune age yielded an R^2^ of 0.899 and MAE of 4.45 years (**Supplemental Table S3E**). This strong correlation demonstrates that global immune age retains accuracy in donors excluded from training of the contributing clocks.

We next examined the elastic-net coefficients to characterize the transcriptional programs contributing to age prediction and confirm that the clocks rely on known immune aging features (**Figure 1F**). T cell models recovered established aging markers, including *LRRN3* and *TSHZ2*.^21, 22^ Genes associated with naïve states, including *IL7R*, *SELL*, *SOX4,* and *BCL11A*, generally had negative coefficients, predominantly in T cell models, consistent with age-related decline of the naïve T cell pool. Conversely, cytotoxic effector genes (*GZMH, GZMK, PRF1* and *GNLY*) had positive coefficients in the memory T cell models, reflecting the accumulation of terminally differentiated effector cells with age.^23^ Interferon-associated genes contributed in a cell type-specific manner; *IRF9* was positive in naïve B, memory B, and monocyte models, whereas *ISG15* was negative in CD4^+^ memory and monocyte models, indicating that interferon signaling changes with age in different directions across cell types. Finally, genes selected in multiple cell types, including the glucocorticoid-induced regulator *TSC22D3* (*GILZ*), the antiproliferative gene *BTG1*, and the stress response markers *S100A6* and *CRIP1*, point to a shared stress and immunoregulatory program associated with aging. Individual coefficients for all clocks are provided in **Supplemental Table S4**. Together, these markers indicate the clocks are built on established immune aging programs, specifically naïve T cell loss and cytotoxic differentiation, coupled with cell type-specific interferon changes and a shared stress response.^14^

### Accelerated immune aging identifies a high-risk phenotype in younger myeloma patients

After developing the immune aging clocks based on ten cell types, we applied them as a unified immune aging metric to identify patients with accelerated immune aging in a clinically annotated scRNA-seq cohort of patients with newly diagnosed multiple myeloma (NDMM) (**Figure 2A**). Patients were analyzed in two age groups, <70 and ≥70 years at diagnosis (**Figure 2B**). Although transplant eligibility is based on fitness rather than age, 70 years is commonly used as an eligibility threshold in clinical trials.^19^ The frequencies of high-risk cytogenetic lesions, including t(4;14), del(17p13), and TP53 mutation, were distributed similarly across age groups, with 1q gain present in 34.5% of patients <70 years old and 44.2% in patients ≥70 years old (**Supplemental Figure 2A**). As expected, younger patients presented with less advanced disease, characterized by lower International Staging System (ISS) stage and a smaller proportion of high-risk IMWG disease. Additionally, this younger cohort more frequently received ASCT and triplet (RVd) induction therapy (**Figure 2C**). Specifically, 59% of patients younger than 70 received ASCT compared with 32% of those 70 or older, and 80% compared to 50% received triplet induction. Although ASCT eligibility is no longer defined by age alone, the 70-year cutoff split this cohort into groups with different treatment distributions, supporting separate analysis of the two age groups.

**Figure 2.**
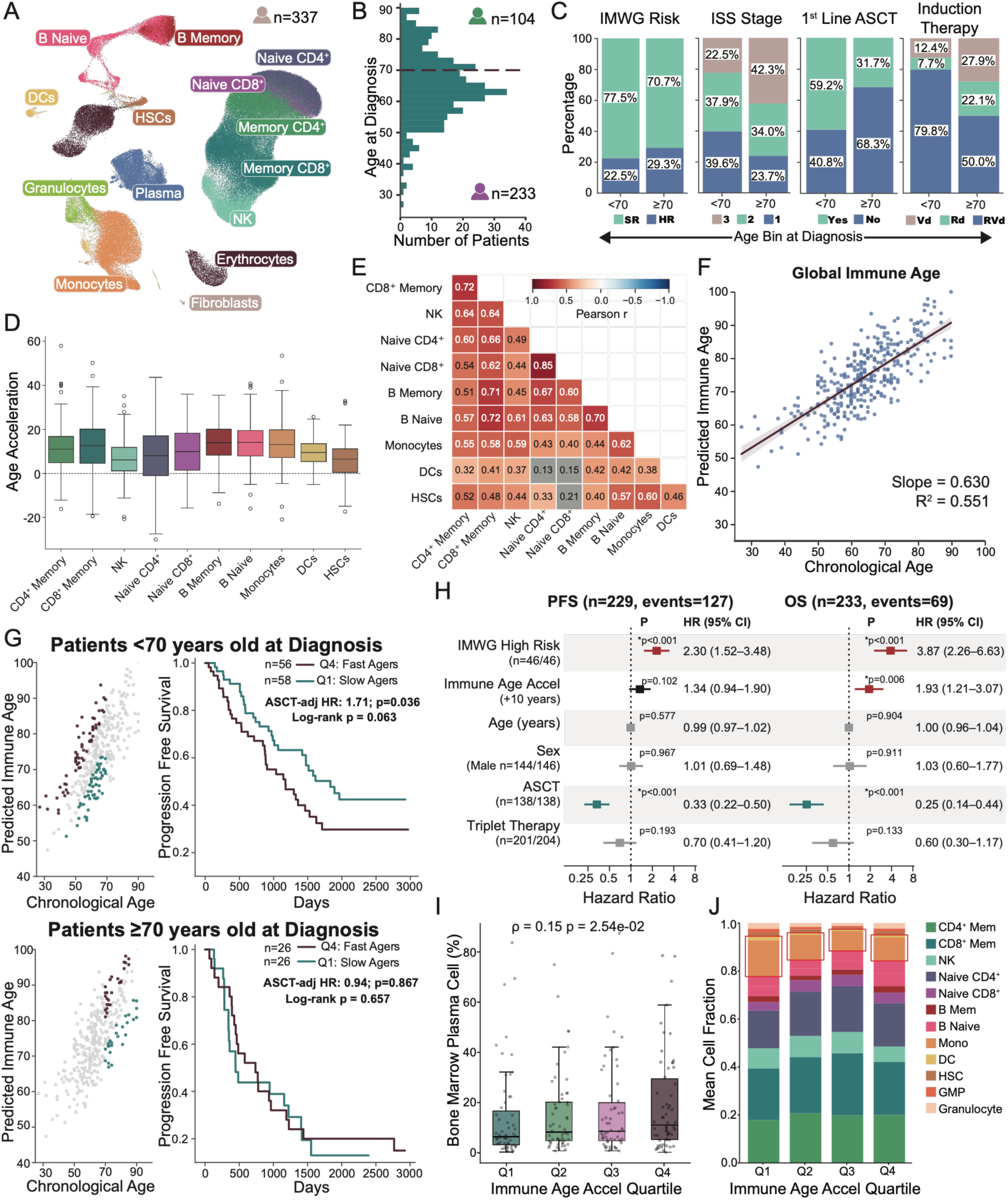
Accelerated immune aging is an independent predictor of survival in multiple myeloma. **(A)** UMAP of MM Immune Atlas (n = 337) with cell subtypes organized into 14 lineages, including the 10 included in the bone marrow immune aging model. **(B)** Histogram of patient age at the time of diagnosis in the Immune Atlas. Dashed line separates the histogram at 70 years of age, with 104 patients 70 years or older and 233 patients under 70. **(C)** Stacked bar plots comparing clinical characteristics of MM patients under 70 and 70 and older at the time of diagnosis. Percentages are of patients with available data. **(D)** Box-and-whisker plot of each cell lineage immune age acceleration in the MM cohort. Boxplots display the median (center line), interquartile range (IQR) (box), whiskers extending to 1.5x the interquartile range, and individual observations beyond whiskers (dots). **(E)** Pairwise Pearson correlation matrix of cell type-specific immune age acceleration across the MM cohort. Color intensity corresponds to the Pearson correlation coefficient (r), with red indicating positive correlations and blue indicating negative correlations. The upper right triangle is omitted for clarity. Non-significant (q ≥ 0.05) correlations are shown in grey. **(F)** Predicted global immune age versus chronological age across patients within the MM cohort. Each point represents one patient. The solid line represents the linear regression fit, and the shaded band denotes the 95% confidence interval (CI). Model performance is summarized by the regression slope and R^2^. **(G)** Association of immune age acceleration with progression-free survival (PFS) in younger and older patients with MM. Patients were stratified into age groups (<70 years and ≥ 70 years), and within each age group, were divided into quartiles based on global immune age acceleration. Left panels show predicted global immune age versus chronological age for all patients (gray), with the respective group’s slowest agers (Q1, teal) and fastest agers (Q4, burgundy) highlighted. Patient numbers reflect quartile members with annotated ASCT and survival data for survival analysis. Right panels show Kaplan-Meier curves for the highlighted patients. Hazard ratios (HRs) were estimated using Cox proportional hazards models adjusted for ASCT status; adjusted HRs, corresponding p-values, and log-rank p-values are displayed. **(H)** Forest plots of multivariable Cox proportional hazards models for PFS and OS in patients <70 years of age. Global immune age acceleration (per 10 years) was adjusted for IMWG high-risk status, chronological age, sex, first-line ASCT, and triplet induction therapy, stratified by collection site. Squares indicate HRs and bars show 95% CIs; n and events are shown for each model. Asterisks indicate p < 0.05 with HR > 1 (red) or HR < 1 (teal). Gray markers are not significant. **(I)** Bone marrow plasma cell burden across age-specific global immune age acceleration quartiles in patients <70 years old. Boxplots display the median (center line), IQR (box), whiskers extending to 1.5x IQR, and individual patients (dots). The association between global age acceleration and plasma cell burden was assessed by Spearman correlation, with ρ and p-value displayed. **(J)** Mean immune cell composition across patients <70 years of age, split into global immune aging quartiles. Cell type abundances were normalized to the total abundance of all immune cell populations for each patient. Stacked bars represent the mean relative cell fraction of each immune cell type within each quartile (Q1-Q4). Red boxes highlight the monocyte and DC fractions.

Applying the cell type-specific clocks to each patient revealed positive median age acceleration values across all ten cell types, supporting that the NDMM cohort exhibits advanced immune aging compared to healthy donors (**Figure 2D**; **Supplemental Figure 2B**). Furthermore, cell type-specific age acceleration scores were positively correlated with one another, suggesting that common microenvironmental influences may contribute to concurrent immune aging along different lineages and supporting their combination into a global score (Pearson r = 0.32-0.85 for significant pairs; **Figure 2E**). Global immune age also positively correlated with chronological age with a slope of 0.630 (R^2^ = 0.551), reflecting that these measures are broadly consistent. However, there are also many patients with considerable deviation between their global immune age and chronological age (**Figure 2F**). To investigate the clinical relevance of this deviation, we next focused our investigation on patients who appeared to demonstrate the highest rates of immune age acceleration. Specifically, we classified patients in the highest quartile of immune age acceleration within their age groups as “fast agers” and compared them to those in the lowest quartile, labeled as “slow agers”. When quartiles are defined by a single-cell type clock, we refer to patients with the most and least accelerated cells of that type.

The assessment of global immune age acceleration in patients younger than age 70 with outcomes depicted that fast agers (Q4, n = 56) had poorer ASCT-adjusted PFS compared to the slow agers (Q1, n = 58; HR = 1.71, p = 0.036; **Figure 2G**). Examining overall survival (OS), fast agers (Q4, n = 58) also had a higher ASCT-adjusted hazard than slow agers (Q1, n = 59; HR = 2.54, p = 0.011, **Supplemental Figure 2C**). These outcome associations were not detected in patients older than 70 years (**Figure 2G**; **Supplemental Figure 2C**; **Supplemental Figure 2G**), and therefore subsequent analyses focused on patients under age 70.

We next investigated whether immune age acceleration was associated with survival independent of established clinical risk factors using multivariable Cox proportional hazards models adjusted for International Myeloma Working Group (IMWG) high-risk status, first-line ASCT, chronological age, sex, and triplet induction therapy, stratified by collection site (**Figure 2H**). Greater global immune age acceleration was significantly associated with shorter OS (HR = 1.93 per 10 years, p = 0.006), while the association with PFS did not reach significance (HR = 1.34, p = 0.102). One standard deviation of global age acceleration corresponded to 7.2 years, with corresponding hazard ratios of 1.6 for OS and 1.23 for PFS (**Supplemental Table S8**).

In contrast, chronological age was not associated with either PFS (HR = 0.99, p = 0.577) or OS (HR = 1.00, p = 0.904) in patients under age 70 using the multivariable models. Furthermore, Kaplan-Meier analysis stratified by decade of age at diagnosis illustrated that chronological age was not associated with PFS (log-rank p = 0.20) or OS (log-rank p = 0.35) in these patients under age 70 years, whereas a significant association between chronological age and both PFS and OS was observed in patients over 70 (log-rank p = 0.005 for OS and PFS; **Supplemental Figure 2D**). Collectively, these findings illustrate that immune age acceleration captures prognostic information beyond chronological age and established clinical risk factors for patients under 70 with MM.

We next assessed whether immune age acceleration reflected tumor burden or differences in BM immune composition by correlating global immune age acceleration with BM plasma cell burden and with the abundance of each immune cell type. Among patients younger than 70 years, global immune age acceleration was weakly correlated with plasma cell burden by flow cytometry (Spearman ρ = 0.15, p = 0.0254), although median tumor burden was significantly higher in fast agers than slow agers (Mann-Whitney U p = 0.033; **Figure 2I**). Similarly, immune composition was largely unrelated to global immune age acceleration (**Figure 2J**; **Supplemental Table S6**), with only monocyte and DC fractions showing weak inverse relations (monocytes ρ = -0.22, p = 0.001; DCs ρ = -0.18, p = 0.006; **Supplemental Figures 2E-F**). Additionally, to confirm that the survival association was not driven by tumor burden or these compositional differences, we added plasma cell burden and the abundance of monocytes and DCs to the multivariable survival models. After adjustment for bone marrow plasma cell burden, global immune age acceleration remained associated with OS (HR = 1.90, p = 0.007), but not PFS (HR = 1.31, p = 0.124; **Supplemental Figure 2I**). In separate models adjusted for monocyte and DC fractions, monocyte abundance was independently associated with OS (HR = 1.51, p = 0.008, **Supplemental Figure 2H**), but not PFS, consistent with the myeloid enrichment previously observed among rapid progressors in this cohort.^18^ Importantly, immune age acceleration remained associated with OS while accounting for these compositional changes (HR = 1.65, p = 0.041). Immune age acceleration therefore reflects transcriptional shifts within BM cell types rather than BM composition, and its association with OS was not explained by tumor burden or by monocyte and DC abundance.

### T and NK cell age acceleration is associated with poor survival independent of NF-κB enrichment, with largest contribution from CD4⁺ memory T cells

To characterize the age-associated transcriptomic changes in T and NK cells, we investigated the pathways associated with predicted immune age in naïve and memory T cells and NK cells in the healthy reference and MM (<70 years old) cohorts, excluding genes used to train the corresponding clocks (**Supplemental Tables S4**, **S5**, and **S10**).

Pathways associated with immune age spanned inflammatory, stress-response, interferon, and metabolic programs (**Figure 3A**; **Supplemental Figure 3A**). TNF-α signaling via NF-κB was the most consistent association, increasing with predicted immune age across T and NK populations in both cohorts. Inflammatory response, hypoxia, apoptosis, and p53 signaling were also positively associated with predicted age, consistent with inflammaging.^24^ In the MM cohort, these associations were also accompanied by increased aggrephagy, the selective degradation of misfolded proteins by autophagosomes, across all five populations, alongside increased autophagy in memory T cells, suggesting additional changes in protein and organelle clearance.^25^

**Figure 3.**
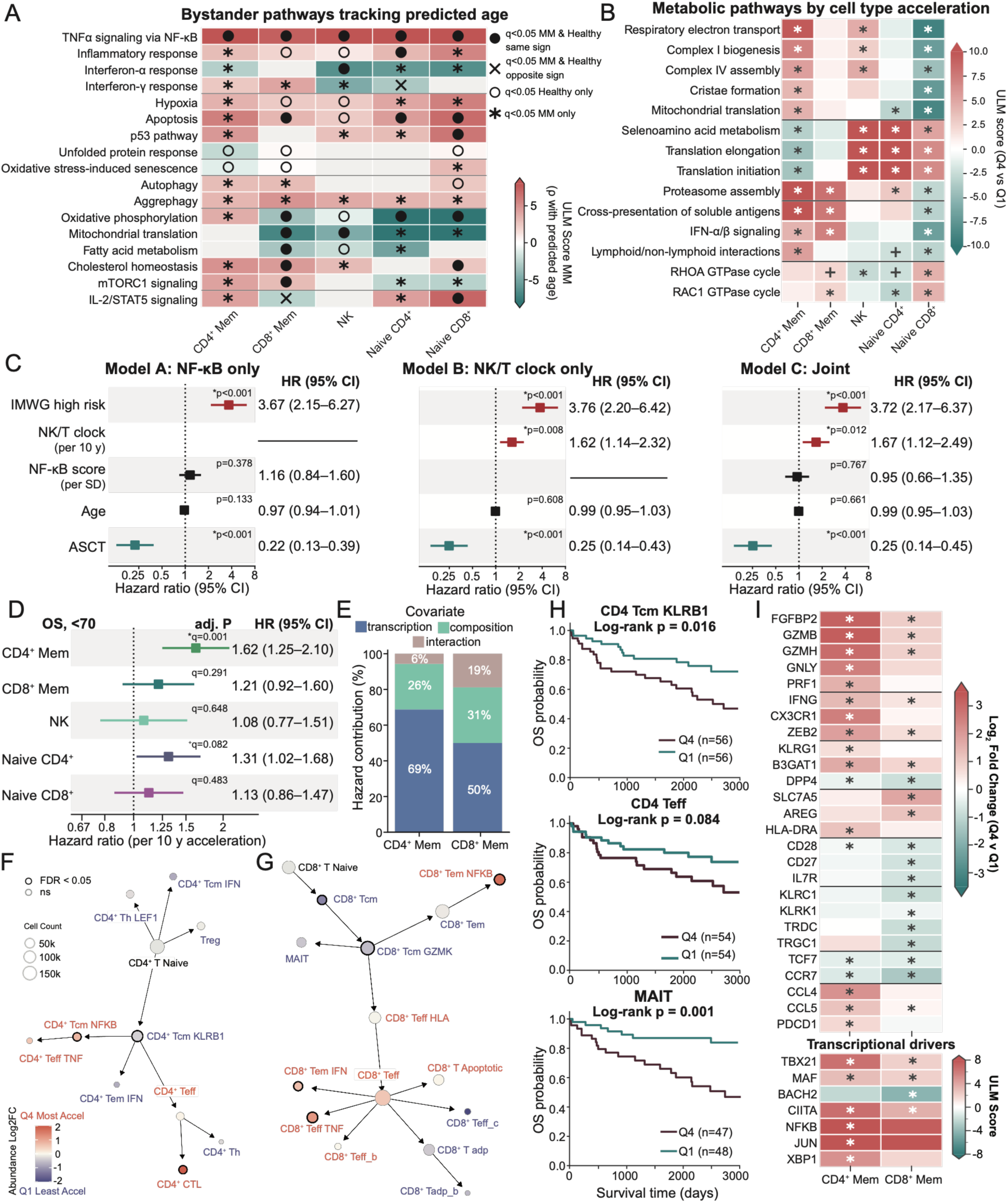
T and NK cell lineage-specific age accelerations are associated with survival independently of NF-κB. **(A)** Pathways associated with predicted age in T and NK lineage clocks in bystander genes (non-clock genes) in bone marrow from healthy and MM cohorts. For each lineage and cohort, genes were ranked by partial Spearman correlation (ρ) with predicted age from the lineage-specific clock (adjusted for collection site, sex, and log median genes per cell; Fisher z-transformed). Pathway activity scores were calculated using a univariate linear model (ULM). Significance is defined as q < 0.05. Color shows ULM score in MM (gray where neither cohort was significant). Symbols indicate significance in cohorts: both MM and healthy in the same direction (filled circle) or opposite direction (×), healthy only (open circle), and MM only (asterisk). Per-cohort scores are shown in **Supplemental Figure 3A**. **(B)** Pathway activity (ULM score) inferred from differential expression in cells of the indicated lineage in patients with the most accelerated (Q4) vs. least accelerated (Q1) cells of that lineage based on the lineage-specific clock in MM patients aged <70 years. Positive scores indicate higher activity in Q4. Asterisks are shown where q < 0.05; plus sign where q < 0.10. **(C)** Forest plots of multivariable Cox models for overall survival (OS) evaluating the NF-κB score alone (Model A), aggregate NK/T clock age acceleration alone (Model B), and both measures together (Model C; n = 233, 69 events). Likelihood ratio tests (LRTs) compare Model C with Model B (NF-κB score added) and Model A (NK/T clock added). Adding NK/T clock age acceleration improved model fit (χ² = 6.25, p = 0.012), whereas adding the NF-κB score did not (χ² = 0.09, p = 0.766). Squares indicate hazard ratios (HRs), and horizontal lines indicate 95% confidence intervals (CIs). Asterisks indicate p < 0.05; significant HRs greater than 1 are shown in red and HRs less than 1 in teal. **(D)** Forest plots of multivariable Cox HRs for OS per 10-year increase in lineage-specific age acceleration in T- and NK-cell lineages. Models were adjusted for chronological age, IMWG high-risk status, sex, triplet therapy, and first-line ASCT, stratified by site. Sample sizes and event counts were: CD4⁺ memory, n = 233 (69 events); CD8⁺ memory, n = 233 (69 events); NK, n = 227 (67 events); naïve CD4⁺, n = 230 (68 events); and naïve CD8⁺, n = 214 (60 events). Q values represent Benjamini–Hochberg (BH)-adjusted p values. Asterisks indicate q < 0.05; plus signs indicate q < 0.1. **(E)** Shift-share decomposition of the OS hazard associated with CD4⁺ and CD8⁺ memory clock acceleration into transcriptional, compositional, and interaction components. Bars show each component’s share of the variance of the Cox linear predictor (Methods). **(F, G)** Subtype abundance shown on subtype differentiation trajectories (see Methods) in patients with the most accelerated versus least accelerated CD4+ memory cells (**F**) and patients with the most accelerated and least accelerated CD8+ memory cells (**G**), defined as top and bottom quartiles of lineage-specific age acceleration. Naïve and regulatory populations are shown for context and were not tested; schematic only: edge length not to scale. Color shows log₂ fold change in subtype proportion (red, enriched in Q4; blue, enriched in Q1); node size shows total cell count. Bold outlines mark subtypes with significant differential abundance (Mann-Whitney U, q < 0.05). **(H)** Kaplan-Meier OS curves comparing patients with the lowest (Q1) and highest (Q4) subtype-level age acceleration in 3 subtypes. Subtype-level age acceleration was obtained by applying the parent clock to the subtype’s pseudobulk expression (CD4⁺ memory clock applied to pseudobulk expression in CD4⁺ Tcm KLRB1 cells and CD4⁺ Teff cells; CD8⁺ memory clock to MAIT cells). **(I)** Differential expression between patients with the most accelerated (Q4) and least accelerated (Q1) CD4⁺ and CD8⁺ memory cells, with quartiles defined by each lineage clock. Top: log₂ fold change of selected genes of interest. Bottom: transcription factor activity scores (ULM, CollecTRI). Asterisk indicates q < 0.05.

Interferon and metabolic pathways exhibited more cell type-specific patterns in MM. Predicted age was associated with weaker interferon-α (IFN-α) response in CD4⁺ memory T, naïve T, and NK cells, and a stronger interferon-γ (IFN-γ) response in memory T cell compartments, consistent with the interferon patterns previously linked to rapid progression in this cohort.^18^ Mitochondrial translation and oxidative phosphorylation generally decreased with predicted age in naïve and CD8^+^ memory T cells, whereas CD4^+^ memory T cells showed increased oxidative phosphorylation (OXPHOS).

To further investigate the contrasting metabolic associations observed between T cells, we compared pathway enrichment between patients with the highest and lowest cell type-specific immune age acceleration (**Figure 3B**). CD4+ memory T cells with greater immune age acceleration showed increased enrichment of mitochondrial respiration pathways and decreased enrichment of translation and selenoamino acid metabolism pathways, whereas naïve T cells showed the opposite pattern, suggesting that CD4^+^ memory cells shift away from protein synthesis and toward oxidative energy production, consistent with previous reports of increased mitochondrial mass and respiratory capacity in senescent CD4^+^ T cells.^26^ In contrast, inflammatory Th1 cells increase glycolysis to support the biosynthetic and proliferative demands of effector activation, while limiting OXPHOS.^27, 28^ The combination of increased mitochondrial and reduced translation-related programs may therefore reflect a metabolic state less supportive of clonal expansion and acute effector function.

Given the consistent association between predicted immune age and NF-κB signaling, we asked whether the clock was simply a readout of inflammatory signaling, which would make it redundant with an NF-κB score. Although NF-κB enrichment was positively correlated with NK/T immune age acceleration (Spearman ρ = 0.59, p < 0.001, **Supplemental Figure 3B**), it was not significantly associated with OS in adjusted Cox models (HR = 1.16, p = 0.378). In contrast, NK/T immune age acceleration was associated with OS both independently (HR = 1.62, p = 0.008) and after adjustment for NF-κB enrichment (HR = 1.67, p = 0.012, **Figure 3C**). Neither measure was significantly associated with PFS (**Supplemental Figure 3C**). These results indicate that the NK/T clocks are not simply a readout of NF-κB inflammatory signaling and carry prognostic information beyond NF-κB-associated signaling.

To determine how individual T and NK cell clocks contribute to the association between global immune age acceleration and OS, we examined each cell type clock separately. Among these populations, CD4^+^ memory T cell age acceleration was associated with OS (HR = 1.62, adj. p = 0.001) and trended with PFS (HR = 1.30, adj. p = 0.062), whereas the remaining clocks were not significantly associated with either outcome (**Figure 3D**; **Supplemental Figure 3D**). Since individual cell type clocks are noisier than the global score, we tested the contribution of each clock directly with leave-one-lineage-out analyses. The OS hazard ratio remained above 1 after each cell type was excluded (**Supplemental Figure 3E**), with the greatest attenuation observed following removal of CD4^+^ memory cells (HR = 1.68, 95% CI 1.06-2.68; **Supplemental Table S12**). These findings suggest that the OS association of the global clock is distributed across cell type models, with CD4^+^ memory cells as the largest single contributor rather than its sole source.

Because memory T cells span multiple differentiation states, we next used shift-share decomposition to assess the relative contributions of transcriptional changes within subtypes, shifts in subtype abundance, and their interaction to the association between immune age acceleration and OS (*see Methods*). Shift-share decomposition attributed 69% of the CD4^+^ memory T cell contribution to OS risk to within-subtype transcriptional differences, with smaller contributions from compositional shifts (26%) and their interaction (5%; **Figure 3E**). In CD8^+^ memory T cells, these contributions were more evenly distributed (50%, 31%, and 19%, respectively). Thus, transcriptional differences within subtypes accounted for the greatest contribution to the modeled OS association in CD4+ memory T cells, whereas transcriptional and compositional contributions were more balanced in CD8+ memory T cells.

We next examined which subtypes account for these compositional shifts by comparing their relative abundance between patients with the highest and lowest cell type-specific age acceleration (**Figure 3F-G**). Patients with the most accelerated CD4^+^ memory T cells had higher abundance of NF-κB^+^ central memory (CD4⁺ Tcm NF-κB) and cytotoxic (CD4⁺ CTL) subtypes, whereas KLRB1^+^ central memory (CD4⁺ Tcm KLRB1) cells were less abundant. Similarly, patients with the most accelerated CD8^+^ memory cells had higher abundance of NF-κB^+^ effector memory (CD8⁺ Tem NF-κB), TNF^+^ effector (CD8⁺ Teff TNF), and type I IFN^+^ effector memory (CD8⁺ Tem IFN) cells, while those with the least accelerated CD8^+^ memory cells had higher abundance of central memory populations (CD8⁺ Tcm and CD8⁺ Tcm GZMK). These differences indicate a shift from central memory toward effector, cytotoxic, and inflammatory subtypes with greater age acceleration. In NK cells, greater age acceleration was similarly associated with differences in subtype composition (**Supplemental Figure 3F**), including reduced CD56_bright_ (adj. p = 0.002) and CD56_dim_ (adj. p < 0.001) fractions and a higher adaptive NK fraction (adj. p < 0.001). Adaptive NK expansion is commonly driven by cytomegalovirus (CMV) infection, but because CMV serology was not available for this cohort, we could not assess its contribution.^29^

Although greater memory T cell age acceleration was associated with shifts toward differentiated effector populations, the significant contribution of within-subtype transcriptional variation prompted us to investigate whether age acceleration within individual memory T cell subtypes also stratified patient survival. Therefore, we applied the CD4^+^ and CD8^+^ memory clocks to their corresponding subtype-level pseudobulk expression profiles and calculated patient-specific age acceleration for each subtype. The CD4⁺ memory clock stratified both OS and PFS when applied to CD4^+^ Tcm KLRB1 cells (log-rank p = 0.016 and p = 0.035, respectively; **Figure 3H; Supplemental Table S13**). Similarly, the CD8⁺ memory clock stratified OS when applied to MAIT cells (log-rank p = 0.0009), although the aggregate CD8⁺ memory profile did not (p = 0.075). Notably, CD4⁺ Tcm KLRB1 cells were less abundant in patients with the most accelerated CD4⁺ memory cells, yet their subtype-specific transcriptional age remained associated with survival. These subtype-level survival associations indicate that the prognostic information captured by the memory T cell clocks reflects aging-associated transcriptional variation within individual subpopulations.

Following these subtype-level survival analyses, we next investigated whether the enrichment of differentiated effector T cell populations among patients with greater immune age acceleration was accompanied by corresponding transcriptional changes. We, therefore, compared gene expression and inferred transcription factor activity between patients with the highest and lowest CD4+ and CD8+ memory T cell age acceleration (**Figure 3I**). Both memory T cell populations showed increased expression of terminal effector- and senescence-associated genes, including *B3GAT1*, *ZEB2*, *GZMB*, and *FGFBP2*, with additional increases in *CX3CR1* and *PDCD1* in accelerated CD4⁺ memory T cells. In contrast, less accelerated cells retained higher expression of co-stimulatory and naïve-like genes, including *CD28*, *TCF7*, and *CCR7*. Consistently, accelerated cells showed higher activity of the positive regulator of terminal differentiation *TBX21* (T-bet) in both memory populations, with lower activity of *BACH2*, a negative regulator of terminal differentiation (**Figure 3I**). Greater age acceleration was also associated with higher levels of stress-response genes and NF-κB targets (**Supplemental Figure 3G**), consistent with our pathway-level analyses. Thus, memory T cell age acceleration appears associated with a senescent-like terminal effector phenotype, alongside a broader NF-κB-associated stress response across T and NK cell compartments.

### B cell, myeloid, and progenitor clocks are linked to inflammatory and metabolic programs, and monocyte acceleration shows a trend toward shorter PFS

Having identified inflammatory and metabolic signatures associated with NK and T cell aging, we next investigated whether these patterns were shared by the remaining monocytes, memory B, naïve B, DC, and HSC populations. Similar to T and NK cells, TNF-α signaling via NF-κB was positively associated with predicted age in monocytes, memory B, naïve B, and DCs in both healthy and MM cohorts (**Figure 4A**; **Supplemental Figure 4A**), identifying TNF-α/NF-κB-associated inflammation as a common feature of predicted immune aging across BM cell types.

**Figure 4.**
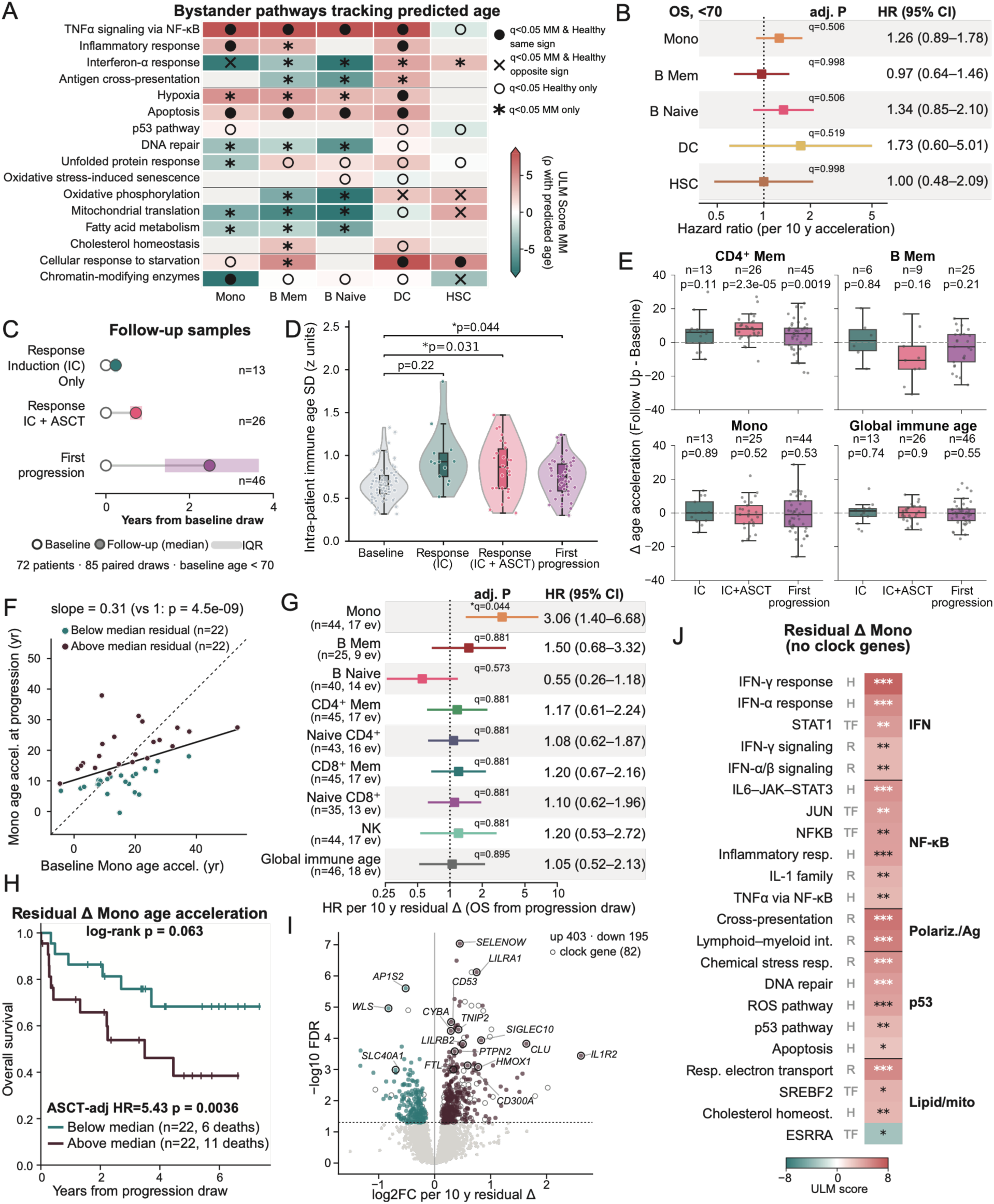
B cell, myeloid, and progenitor clocks are associated with inflammatory and metabolic programs, and increased monocyte age acceleration at first progression is associated with subsequent overall survival. **(A)** Pathways associated with predicted age in monocyte, memory B, naïve B, DC, and HSC clocks in bystander genes in bone marrow from healthy and MM cohorts. For each lineage and cohort, genes were ranked by partial Spearman correlation (ρ) with predicted age from the lineage-specific clock (adjusted for collection site, sex, and log median genes per cell; Fisher z-transformed). Pathway activity scores were calculated using a univariate linear model (ULM). Significance is defined as q < 0.05. Color shows ULM score in MM (gray where neither cohort was significant). Symbols indicate significance in cohorts: both MM and healthy in the same direction (filled circle) or opposite direction (x), healthy only (open circle), and MM only (asterisk). Per-cohort scores are shown in **Supplemental Figure 4A**. **(B)** Forest plots of multivariable Cox HRs for OS per 10-year increase in lineage-specific age acceleration in MM patients younger than 70 years. Models were adjusted for chronological age, IMWG high-risk status, sex, triplet therapy, and first-line ASCT, stratified by site. Q values represent BH-adjusted p values. Sample sizes and event counts were: monocytes, n = 228 (65 events); memory B cells, n = 197 (56 events); naïve B cells, n = 207 (54 events); dendritic cells, n = 104 (25 events); and HSCs, n = 94 (29 events). Asterisks indicate q < 0.05; plus signs indicate q < 0.1. **(C)** Timing of paired follow-up samples in 72 MM patients aged <70 years at baseline (85 paired draws). Follow-up samples were taken after response to induction chemotherapy alone (IC; n = 13), after response to induction plus ASCT (IC + ASCT; n = 26), or at first progression (n = 46). Open circles mark baseline, filled circles indicate the median follow-up time. Shaded bars span IQR. **(D)** Within-patient standard deviation (SD) of z-transformed lineage age accelerations at baseline and in each follow-up group. Accelerations for 8 lineages (excluding DC and HSC due to low cell numbers) were z-scored against the baseline mean and SD of patients with follow-up samples. All samples with ≥4 lineages are shown. p-values are from paired Wilcoxon signed-rank tests of each follow-up group against the same patients’ baseline samples for patients with ≥4 lineages in both baseline and follow-up samples. Dots show individual samples, violins show SD distribution, and underlying boxplots show median (line), IQR (box), and 1.5x IQR (whiskers) in z units. **(E)** Change in age acceleration from baseline (follow-up − baseline) for CD4⁺ memory, memory B, monocyte, and global immune age, by follow-up group. The n and p-values are from Wilcoxon signed-rank tests of Δ against zero. Boxplots show the median (line), interquartile range (box), and 1.5× IQR (whiskers) in acceleration years. Other lineages are shown in **Supplemental Figure 4E**. **(F)** Monocyte age acceleration at first progression versus baseline monocyte age acceleration. The line shows the linear fit of progression acceleration on baseline acceleration. Points are colored by residualized Δ above (burgundy) or below (teal) the median. **(G)** Forest plot of OS HRs after the first progression draw, per 10 years of residualized Δ age acceleration for each lineage clock and for global immune age. Cox models were adjusted for chronological age (per 10 years), sex, IMWG risk, and first-line ASCT, stratified by site, and fitted with a ridge penalty (λ = 0.1). The number of patients and events is shown for each model; asterisk indicates q < 0.05. **(H)** Kaplan-Meier curves of OS after progression draw, with patients split at the median residualized Δ monocyte age acceleration. Patients with high residualized Δ monocyte age acceleration are shown in burgundy, low in teal. Tick marks show censored patients. **(I)** Volcano plot of genes associated with residualized Δ monocyte age acceleration in pseudobulk monocyte profiles from first-progression samples (DESeq2; design ∼ collection site + sex + log median genes per cell + residualized Δ). The x-axis shows log₂ fold change per 10 years of residualized Δ and the y-axis shows −log₁₀ false discovery rate (FDR). The dotted line marks FDR = 0.05. Colored points are significantly upregulated (burgundy, n = 403) or downregulated (teal, n = 195) genes. Open circles mark significant clock genes (n = 82). **(J)** Pathway and transcription factor activity (ULM score) associated with residualized Δ monocyte age acceleration, excluding clock genes. Gene sets are Hallmark (H), Reactome (R), and CollecTRI transcription factor (TF) regulons, grouped into interferon, NF-κB, polarization/antigen presentation, p53, and lipid/mitochondrial programs. *q < 0.05, **q < 0.01, ***q < 0.001.

In contrast, interferon and antigen-presentation pathways showed cell type-specific associations. In MM, IFN-α response and antigen cross-presentation were negatively associated with predicted age in B cells but positively associated in DCs. Monocytes also showed opposing IFN-α response between cohorts with negative associations with predicted age in MM and positive associations in healthy donors. These differences highlight the heterogeneity of age-associated interferon signaling across immune populations and clinical contexts.

Beyond inflammatory signaling, predicted age was associated with distinct stress-response and metabolic changes, particularly in monocytes and B cells. Apoptosis and hypoxia were positively associated with predicted age in monocytes, DCs, and B cells, while DNA repair, mitochondrial translation, and fatty acid metabolism were negatively associated with predicted age in monocytes and both B cell populations in MM, together with reduced unfolded protein response in monocytes and oxidative phosphorylation in B cells. Together, these associations suggest that older-appearing monocytes and B cells in MM exhibit reduced DNA repair and mitochondrial metabolic activity.

We next evaluated whether these pathway-level associations were reflected in the gene expression differences between patients with the highest and lowest cell type-specific age acceleration (**Supplemental Figure 4D**). Monocytes from the most accelerated group showed increased expression of *NAMPT*, an inflammation-induced enzyme of NAD⁺ metabolism, and *PLIN2*, a lipid-droplet protein that marks lipid-laden inflammatory myeloid cells, along with the interferon-stimulated genes *IRF1*, *IFITM2*, *LY6E*, and *ISG15*. Interferon-stimulated genes were also higher in the most accelerated DCs. In B cells, greater age acceleration was associated with increased expression of the pro-apoptotic genes *BBC3* and *BCL2L11* and reduced expression of the innate sensing and signaling genes *TLR10* and *TRAF3IP3*. These profiles highlight inflammatory and lipid-associated changes in accelerated monocytes, increased interferon-responsive signatures in accelerated monocytes and DCs, and pro-apoptotic signaling with reduced innate sensing in accelerated B cells.

Finally, to assess their prognostic relevance, we tested whether age acceleration in these cell types was independently associated with survival using multivariable Cox models. None of the individual cell type clocks were significantly associated with OS (**Figure 4B**), although monocyte age acceleration showed a trend toward shorter PFS (HR = 1.41, adj. p = 0.072; **Supplemental Figure 4B**) without significant differences in monocyte subset composition between the most and least accelerated groups (all adj. p > 0.66; **Supplemental Figure 4C**). Overall, the B cell, monocyte, DC, and HSC clocks capture distinct age-associated inflammatory and metabolic changes but show limited evidence of prognostic value individually.

### Baseline-adjusted change in monocyte age acceleration at progression is associated with outcomes

To ascertain whether immune age acceleration changes with treatment or at progression, and whether these changes carry prognostic information beyond baseline, we examined within-patient changes in immune age acceleration during follow-up. Longitudinal samples were available for 72 patients younger than 70 years at diagnosis, comprising 85 follow-up samples with paired baseline draws. Of these, follow-up samples were collected after induction chemotherapy alone (IC, n = 13), induction chemotherapy followed by ASCT administration (IC + ASCT, n = 26), or at first recorded progression (n = 46), with a median interval of 2.48 years from baseline to first progression (**Figure 4C**). Compared with baseline samples, the within-patient standard deviation among cell type-specific age acceleration scores was higher after IC + ASCT (p = 0.031) and at first progression (p = 0.044), but not after IC alone (p = 0.22; **Figure 4D**). This increased variability following ASCT and at progression suggested that longitudinal changes in immune aging may significantly differ between cell types, prompting us to examine cell type clocks individually in follow-up samples.

Changes in age acceleration from baseline (Δ) varied by compartment and follow-up group (**Figure 4E**; **Supplemental Figure 4E**). CD4^+^ memory T cell acceleration increased after IC + ASCT (p = 2.3 × 10^-5^) and at first progression (p = 0.0019), but not after IC alone (p = 0.11), whereas naïve CD4⁺ acceleration decreased at first progression (p = 9.2 × 10^-4^). In contrast, global immune age acceleration did not change across any follow-up group. This increased CD4^+^ memory T cell age acceleration after IC + ASCT and at first progression is consistent with shifts toward the cytotoxic, terminally differentiated state observed in accelerated CD4^+^ memory cells at baseline, while naïve CD4^+^ cells moved in the opposite direction at progression, highlighting how a global immune age score may obscure distinct aging trajectories across T cell populations.

Unlike CD4^+^ memory and naïve CD4^+^ T cell age acceleration, which shifted consistently at first progression, monocyte age acceleration showed no net change among follow-up timepoints relative to baseline but varied widely between patients (median Δ = -1.0, IQR = 15.3, **Figure 4E**). Because this heterogeneity would be missed by global immune age comparisons, we next examined patient-level age-associated monocyte changes at follow-up. Monocyte age acceleration at first progression was moderately correlated with baseline (r = 0.46, p = 1.6 × 10^-3^), although the regression slope was significantly below 1 (β = 0.31, p = 4.5 × 10^-9^; **Figure 4F**), indicating regression toward the mean. As a result, the raw Δ monocyte age acceleration mostly reflected baseline acceleration (**Supplemental Figure 4G**). Therefore, to remove this dependence, we adjusted each patient’s Δ monocyte age acceleration for baseline acceleration and time between draws. The resulting residualized Δ captures changes in monocyte age acceleration beyond that expected from baseline and follow-up duration (**Supplemental Figure 4H**).

Greater residualized Δ monocyte acceleration at first progression was associated with OS in a penalized multivariable model adjusted for age at baseline, sex, IMWG high risk status, and first-line ASCT, stratified by site (HR = 3.06, p = 0.005, q = 0.044, n = 44, 17 events; **Figure 4G**), as was raw Δ monocyte acceleration at first progression (HR = 1.83, p = 0.015; **Supplemental Figure 4F**). No other cell type-specific age acceleration measure, nor the global immune age, was associated with OS following the first progression. Similarly, in a median-split Kaplan-Meier analysis, patients with greater residualized Δ monocyte acceleration had a higher ASCT-adjusted HR for OS following progression (HR = 5.43, p = 0.0036; log-rank p = 0.063, **Figure 4H**). While sample size is limited, these findings offer initial evidence that changes in monocyte-specific immune aging may capture prognostic information at progression not reflected by the global immune age score.

To characterize the monocyte state associated with greater age acceleration at progression, we next examined gene expression and pathway associations with residualized Δ monocyte age acceleration. Residual Δ monocyte acceleration was associated with 403 significantly upregulated and 195 significantly downregulated genes (FDR < 0.05, **Figure 4I**), including elevated expression of inhibitory receptors (*LILRB2, CD300A,* and *SIGLEC10*), stress response genes (*CLU* and *HMOX1*), and the IL-1 decoy receptor *IL1R2*. Pathway analysis excluding clock features identified associations with interferon response programs (IFN-γ, IFN-α, STAT1), inflammatory signaling (IL-6–JAK–STAT3, JUN, NFKB, IL-1 family, TNF-α via NF-κB), oxidative stress, DNA repair, and apoptosis, alongside lower ESRRA activity, a regulator of mitochondrial metabolism (**Figure 4J**). Between baseline and progression, changes in NF-κB, hypoxia, ROS, p53, and apoptosis differed between patients above and below the median residualized Δ monocyte acceleration, whereas changes in interferon programs did not (**Supplemental Figure 4I**). Thus, greater-than-expected monocyte aging at progression appears to be accompanied by an inflammatory and immunoregulatory state.

### Fast agers show convergent TNF, CCL3 and IFN-γ signaling from memory lymphocytes and NK cells, with NF-κB activation in MM cells

Building on the age-associated transcriptional programs identified within individual immune populations, we next investigated how communication between cells differs between fast and slow agers. We compared the ligand-receptor interactions in the BM of patients <70 with the highest (fast agers, Q4) and lowest (slow agers, Q1) global immune age acceleration.

Consistent with the TNF-α/NF-κB enrichment observed across immune populations, TNF was one of the strongest communication pathways in fast agers. TNF signaling through TNFR1 (*TNFRSF1A*) and TNFR2 (*TNFRSF1B*) was identified between memory lymphocytes and multiple receiving populations, including monocytes, DCs, NK cells, granulocytes, and other memory cells, in fast agers, whereas no TNF interactions passed the specificity threshold in slow agers (p < 0.05; **Figure 5A**). These differences were driven by higher *TNF* expression in CD8^+^ memory T cells (log_2_fold-change (log_2_FC) = 0.581, adj. p = 0.017), CD4^+^ memory T cells (log_2_FC = 0.58, adj. p = 0.017), and memory B cells (log_2_FC = 1.31, adj. p = 0.001), rather than changes in receptor expression (**Supplemental Figure 5A-B**). Furthermore, *TNF* expression in these senders was correlated with increased immune age acceleration (Spearman ρ = 0.25-0.43, p < 0.001, **Figure 5B**), accompanied by increased TNF-α/NF-κB pathway scores (Spearman ρ = 0.42-0.63, p<0.001) and NFKB1/RELA regulon activity across seven receiving populations (Spearman ρ = 0.39-0.52, ρ = 0.39-0.57, respectively, p < 0.001; **Figure 5C**, **Supplemental Figure 5C**). Together, these results reveal that the BMME in fast agers is characterized by increased TNF production from memory lymphocytes and widespread NF-κB activation in TNF-stimulated immune populations across compartments.

**Figure 5.**
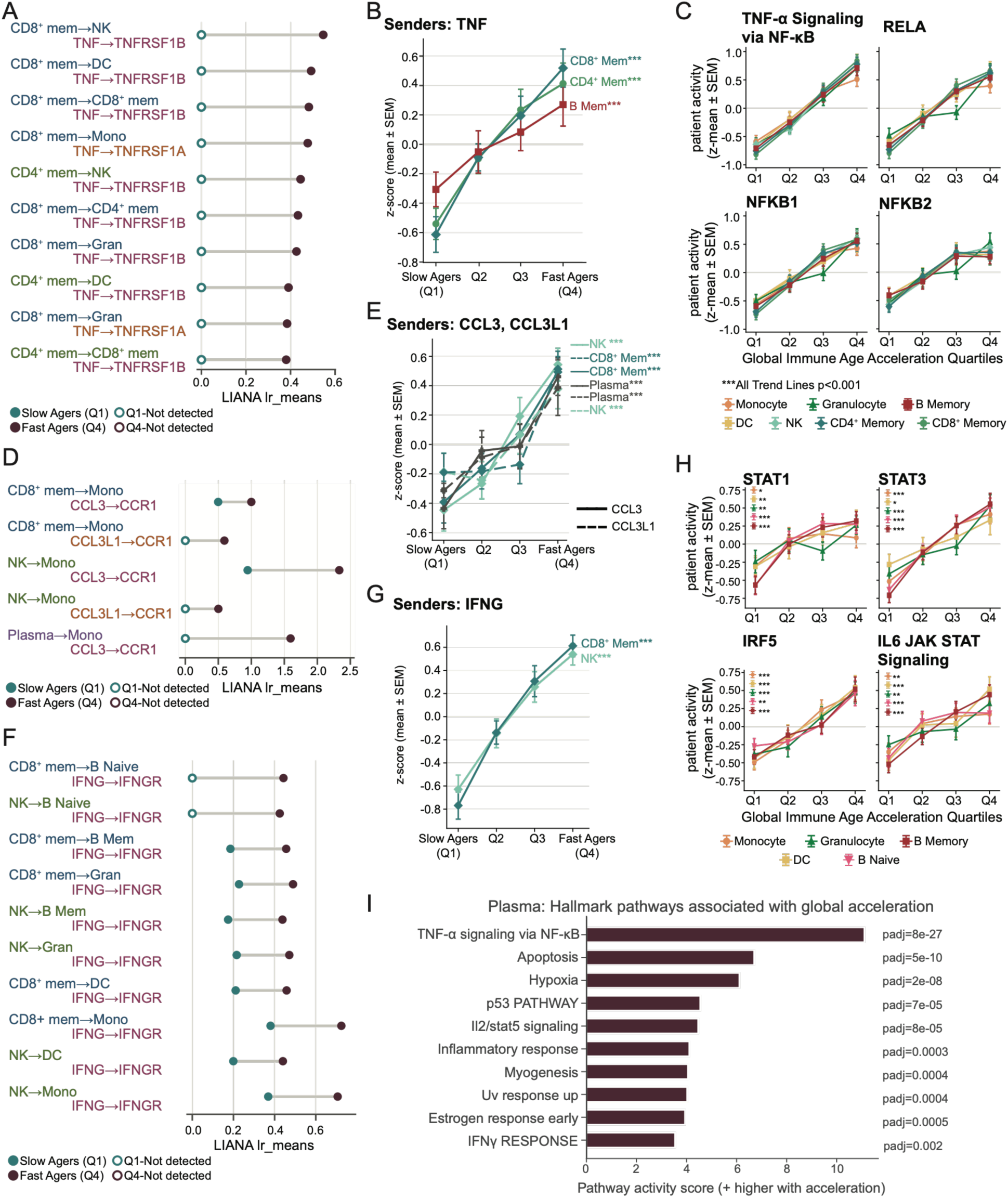
Fast immune agers gain TNF, CCL3/CCL3L1 and IFN-γ signaling from memory lymphocytes and NK cells, with matching downstream responses in the receiving cells. **(A)** Top 10 TNF–TNFRSF1A and TNF–TNFRSF1B ligand–receptor pairs ranked by absolute change in mean interaction score between Q4 and Q1 (LIANA *lr_means*). Rows represent sender–receiver cell-type pairs, color-coded by receptor type: TNFRSF1A (*reddish-brown text*) or TNFRSF1B (*deep magenta text*). Senders are restricted to CD4+ memory (*olive green text*) and CD8+ memory (*deep slate blue text*) T cells. Dot positions indicate interaction strength (*lr_means*) in Q1 (*teal*) and Q4 (*burgundy*). Dot styling denotes significance via CellPhoneDB: filled circles, p < 0.05; open gray circles, p > 0.05; open colored circles at 0, undetected due to low cell-type expression thresholds. **(B)** Stepwise increase of TNF expression with immune age in memory lymphocyte subsets. Line plots show TNF expression trajectories across immune age quartiles (Q1–Q4) for key sender cell types (colored by cell type, CD8+ memory: *deep teal*, CD4+ memory: *sage green*, B memory: *dark red*). Expression values represent patient-level pseudobulk log2CPM, z-score normalized across individuals within each cell type; points and error bars depict mean ±SEM per quartile. Asterisks adjacent to cell-type labels denote the significance of monotonic trends across quartiles (Spearman rank correlation; *p < 0.05, **p < 0.01, ***p < 0.001). **(C)** Downstream signaling and transcriptional activity across immune age quartiles (Q1–Q4) for designated TNF-receiver cell types. Signatures encompass the MSigDB Hallmark TNFα Signaling via NF-κB gene set, and CollecTRI regulons for NF-κB subunits (NFKB1, RELA, NFKB2). Activity scores were computed per single cell using decoupleR univariate linear modeling (ulm), averaged per donor, and z-score normalized across donors within each cell type; error bars indicate mean ± SEM per quartile. Asterisks in the legend denote the significance of monotonic trends across quartiles determined by Spearman rank correlation with ***p < 0.001. **(D)** Ligand-receptor interactions between CCL3 (*deep magenta text*) or CCL3L1 (*reddish-brown text*) and CCR1 across immune age quartiles. Senders are restricted to CD8+ memory (*deep slate blue text)*, NK (*olive green text*), and plasma (*deep violet*) cells. Dot positions indicate interaction strength (lr_means) in Q1 (teal) and Q4 (burgundy). Dot styling denotes significance via CellPhoneDB: filled circles, p < 0.05; open gray circles, p > 0.05; open-colored circles at 0, undetected due to low cell-type expression thresholds. **(E)** CCL3 (solid lines) and CCL3L1 (dashed lines) expression trajectories across immune age quartiles in key sender subsets (NK, memory CD8+ T, and plasma cells). Expression values represent patient-level pseudobulk log2CPM, z-score normalized across individuals within each cell type; points and error bars depict mean ± SEM per quartile. Asterisks adjacent to cell-type labels denote the significance of monotonic trends across quartiles (Spearman rank correlation; *p < 0.05, **p < 0.01, ***p < 0.001). **(F)** Ligand–receptor interactions between IFNG and the heterodimeric IFNGR1/IFNGR2 receptor complex across immune age quartiles. Senders are restricted to CD8+ memory (*deep slate blue text*) and NK (*olive green text*) cells. Dot positions indicate interaction strength (*lr_means*) in Q1 (*teal*) and Q4 (*burgundy*). Dot styling denotes significance via CellPhoneDB: filled circles, p < 0.05; open gray circles, p > 0.05; open-colored circles at 0, undetected due to low cell-type expression thresholds. **(G)** Line plots show *IFNG* expression trajectories across immune age quartiles (Q1–Q4) for key sender cell types (colored by cell type). Expression values represent patient-level pseudobulk log2CPM, z-score normalized across individuals within each cell type; points and error bars depict mean ± SEM per quartile. Asterisks adjacent to cell-type labels denote the significance of monotonic trends across quartiles (Spearman rank correlation; *p < 0.05, **p < 0.01, ***p < 0.001). **(H)** Downstream signaling and transcriptional activity across immune age quartiles (Q1–Q4) for designated IFN-receiver cell types. Regulon activity score for STAT1, alongside downstream STAT3 and IRF5 regulons and Hallmark IL-6/JAK/STAT3 pathway activity. **(I)** Hallmark pathways enriched in plasma associated with global acceleration. Horizontal bar plot showing the top biological pathways associated with global acceleration in plasma cells, ranked by pathway activity score (x-axis; higher values indicate increased activity with acceleration). Adjusted p-values (padj) are displayed to the right of each corresponding bar.

Given the strong associations of monocyte abundance and longitudinal age acceleration with OS, we next investigated whether monocytes received altered intercellular communication in fast agers. Chemokine signaling through the CCR1 receptor via CCL3 (MIP-1α) and CCL3L1 was concentrated almost exclusively on monocytes and was stronger in fast agers, particularly from NK and CD8+ memory T cell populations (**Figure 5D**). Similar to TNF, these differences were associated with increased ligand expression rather than changes in monocyte *CCR1* expression. Expression of *CCL3* was upregulated in NK, CD8+ memory T, and plasma cells (log_2_FC = 0.33-1.63, p < 0.025; **Supplemental Figure 5D**) and significantly correlated with increased immune aging (Spearman ρ = 0.32-0.39, p < 0.001; **Figure 5E**). Given the established role of CCL3-CCR1 signaling in osteoclast differentiation,^30^ these interactions may also involve osteoclast populations that are poorly represented in BM aspirate-based single-cell datasets. In monocytes, inflammatory response, hypoxia, and apoptosis scores also increased markedly with immune age acceleration, with smaller increases in IL-6/JAK/STAT3 signaling and increased activation in IRF5, HIF1A, and NFKB1 (**Supplemental Figure 5E**). These results are consistent with the recruitment and activation of monocytes via CCL3/CCL3L1 ligands from cytotoxic lymphocytes in fast agers, promoting an inflammatory, M1-like state.

Aligning with the increase in IFN-γ with predicted age in memory T cells, we next sought to examine the IFN-γ communication axis. Inferred IFN-γ signaling to IFNGR1 and IFNGR2 receptors on B cell and myeloid lineages was approximately twofold stronger in fast agers (**Figure 5F**). This increase was driven mainly by higher *IFNG* expression in memory CD8^+^ T cells and NK cells, which increased progressively with age acceleration (Spearman ρ = 0.45-0.52, p < 0.001, **Figure 5G**), while receptor expression remained largely unchanged. Receiving populations showed increased expression of canonical STAT1 activity alongside STAT3 and IRF5 regulon activity (**Figure 5H**), with the increased IFN-γ response persisting after excluding genes shared with the IFN-α response signature (**Supplemental Figure 5G**). Notably, although the IFN-γ response within NK cells decreased with predicted age (**Figure 3A**), *IFNG* expression in NK cells was higher in fast agers, highlighting a potential age-associated shift toward adaptive NK cells, which are strong IFN-γ producers.^18^ In summary, IFN-γ signaling from cytotoxic lymphocytes was increased in BM of fast agers, consistent with the shift toward type II interferon stimulation observed in memory T cells.

The widespread inflammatory signaling associated with accelerated immune aging raised the possibility that malignant plasma cells within the same BMME might exhibit corresponding transcriptional changes. Although the single-cell cohort was generated from CD138-depleted BM aspirates, residual plasma cells were retained due to high tumor burden and were previously shown to carry patient-specific copy-number alterations consistent with malignancy.^18^ We therefore examined associations between immune age acceleration and malignant plasma cell gene expression, observing that global immune age acceleration was associated with the differential expression of 21 genes, including the upregulation of cytoskeletal and adhesion (*LMNA*, *CORO1C*, *EZR*, *RHOB*) and glucocorticoid response genes (*FKBP5*, *PER1*, *GLUL*; FDR < 0.05; **Supplemental Figure 5I**). At the pathway level, fast agers also exhibited increased TNF-α signaling via NF-κB, along with increased apoptosis, hypoxia, p53, IL-2/STAT5, and IFN-γ response signatures in their plasma cells (**Figure 5I**). Global immune age acceleration was not associated with t(4;14), t(11;14), t(14;16), del(17p), or 1q gain (Kruskal-Wallis p = 0.051-0.32; **Supplemental Figure 5H**), although TP53 mutation was associated with lower acceleration after adjustment for sample processing site (p = 0.012, q = 0.12, n = 14), suggesting immune aging is largely independent of broad tumor genetics. Together, these results demonstrate that the pro-inflammatory BMME of fast agers extended to NF-κB activation in malignant plasma cells, a pathway linked to MM cell survival.^31, 32^

Across all three signaling axes, differences between fast and slow agers were primarily associated with increased ligand expression in memory lymphocytes and NK cells rather than changes in receptor expression. Inferred TNF, CCL3/CCL3L1, and IFN-γ signaling converged on NF-κB and STAT activation in receiving cells, with NF-κB activation extended to malignant plasma cells. Collectively, these results support that fast agers have a BMME shaped by chronic, lymphocyte-driven inflammatory signaling and corresponding transcriptional changes across immune and tumor populations.

### Immune age remains independently prognostic when modeled with ECOG performance status and laboratory deficits in patients under 70 at diagnosis

To determine whether immune age acceleration captures prognostic information beyond established measures of patient fitness, we compared it with ECOG performance status and a five-item laboratory deficit score (LDS-5). ECOG performance status was available for 298 of 337 patients and was associated with both OS and PFS in patients younger and older than 70 years (ordinal Cox < 0.02 for all comparisons; **Figure 6A**; **Supplemental Figure 6A**). Kaplan-Meier analyses showed that higher ECOG scores were generally associated with worse survival, although individual category comparisons varied by age group. In contrast, LDS-5, available for 334 patients with a mean score of 1.39 deficits, showed more age-dependent prognostic associations (**Figure 6B**). Among patients younger than 70, higher LDS-5 scores were associated with shorter OS (ordinal Cox p = 1.6 × 10^-4^), with a weaker association for PFS (ordinal Cox p = 0.041, **Figure 6C**). Neither association was statistically significant in patients aged greater than 70 years (**Figure 6C**; **Supplemental Figure 6B**). Thus, while ECOG performance status was prognostic across both age groups, LDS-5 was mainly prognostic for OS in younger patients.

**Figure 6.**
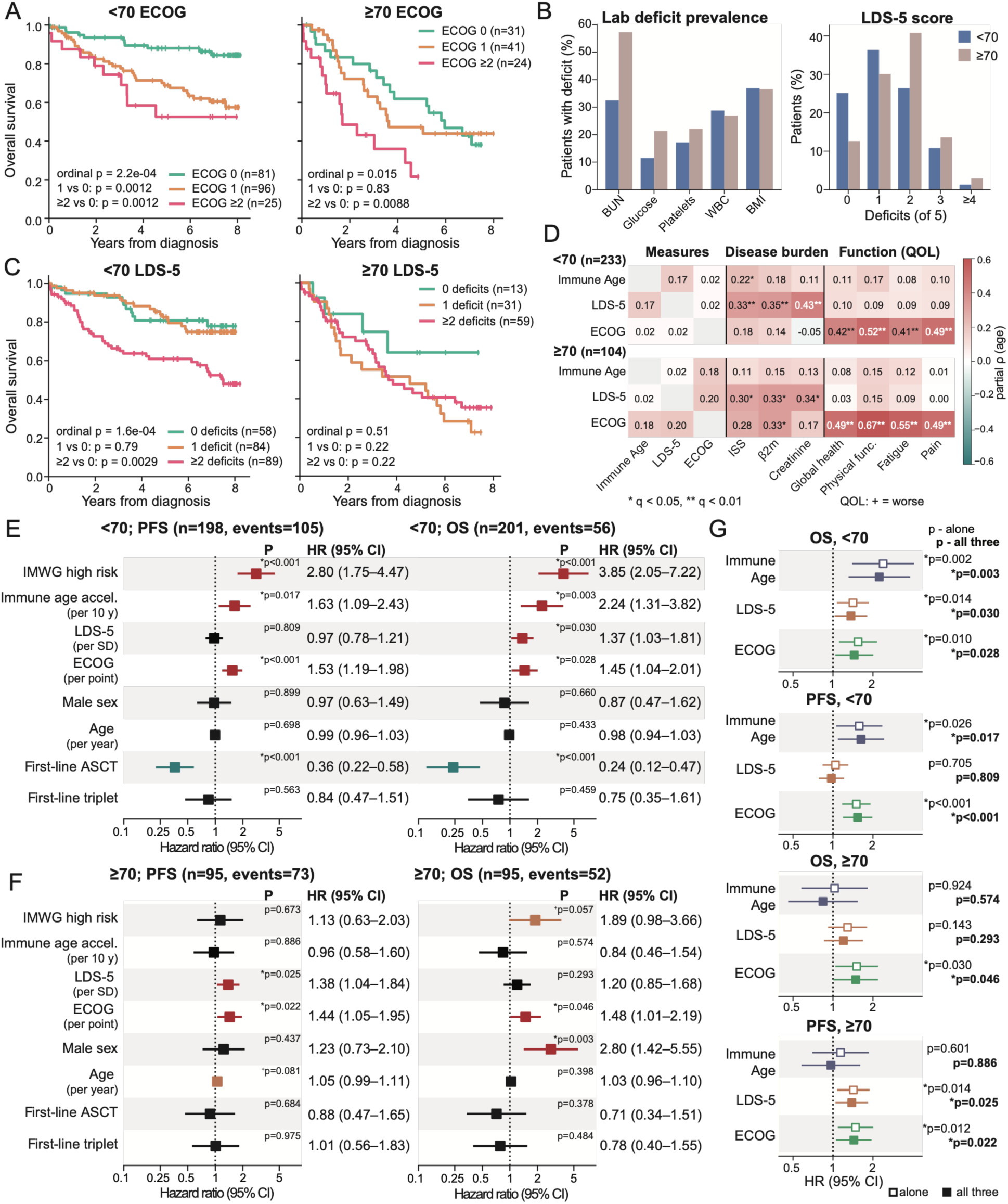
Immune age acceleration, ECOG performance status, and laboratory deficits capture independent dimensions of patient fitness. **(A)** Kaplan-Meier curves showing overall survival (OS) in patients <70 and ≥70 years of age at diagnosis, stratified by ECOG performance status (0 in teal, 1 in orange, ≥2 in pink). Ordinal p-values are from univariable Cox models with ECOG as an ordinal term (likelihood-ratio test). Pairwise comparisons with ECOG 0 used log-rank tests with Holm correction. Tick marks indicate censored patients; curves are truncated when fewer than 3 patients remain at risk. **(B)** Deficit prevalence and score distribution for the five-item laboratory deficit score (LDS-5). Left, prevalence of each deficit by age group. Right, distribution of LDS-5 scores by age group. Scores were computed for patients with at least 4 of 5 items available, scaled to 5 when one was missing (rounded to integers for display). **(C)** Kaplan-Meier curves of overall survival by LDS-5 (0 deficits in teal, 1 in orange, ≥2 in pink) in patients <70 and ≥70 years of age, with statistics as in (A). **(D)** Age-adjusted partial Spearman correlations of global immune age acceleration, LDS-5, and ECOG with each other, with disease burden (ISS stage, β2-microglobulin, creatinine), and with baseline quality-of-life scales (global health, physical functioning, fatigue, pain), shown separately for patients <70 and ≥70 years. Quality-of-life scales are oriented so that positive values indicate worse quality of life. p-values were corrected by the Benjamini–Hochberg method within each age group; * q < 0.05, ** q < 0.01. **(E, F)** Forest plots of multivariable Cox models for PFS and OS in patients <70 (**E**) and ≥70 (**F**) years of age. Each model includes global immune age acceleration (per 10 years), LDS-5 (per standard deviation), and ECOG (per point), adjusted for IMWG high-risk status, chronological age, sex, first-line ASCT, and first-line triplet induction; analyses are stratified by collection site. Analyses include patients with complete data. Squares indicate hazard ratios and bars indicate 95% confidence intervals, shown to the right. Asterisks indicate p < 0.05 with HR > 1 (red) or HR < 1 (teal); plus signs denote p < 0.1 (orange). **(G)** Hazard ratios for immune age acceleration, LDS-5, and ECOG, each modeled alone (open squares, p-values in non-bold font) or with all three measures together (filled squares, p-values shown in bold), for OS and PFS in each age group. All models include the covariates and collection-site stratification described above. Markers as described above.

We next compared immune age acceleration, ECOG, and LDS-5 with measures of disease burden and quality-of-life (QOL) using Spearman correlations adjusted for chronological age to assess whether immune age acceleration represents a distinct measure of patient fitness (**Figure 6D**; **Supplemental Figure 6C**). Among patients under 70 years old, immune age acceleration was not correlated with ECOG (ρ = 0.02, q = 0.88) or LDS-5 (ρ = 0.17, q = 0.056) and modestly associated with ISS stage (ρ = 0.22, q = 0.016) but not significantly correlated with β2-microglobulin or creatinine. In contrast, LDS-5 was correlated with ISS, β2-microglobulin, and creatinine, while only ECOG was associated with all four QOL measures collected using the European Organization for Research and Treatment of Cancer (EORTC) questionnaire. Among patients 70 or older, immune age acceleration was not correlated with any of these measures, whereas associations of LDS-5 with disease burden and of ECOG with QOL persisted. Similarly, only a few cell type-specific associations with disease burden reached significance (q < 0.05, **Supplemental Figure 6C**). Immune age acceleration was therefore largely independent of functional status, QOL, and laboratory deficits in both age groups and only modestly related to disease stage in younger patients.

To assess whether the three measures provided independent prognostic information, we included all three into Cox models adjusted for age, sex, IMWG risk, first-line ASCT, and first-line triplet therapy, stratified by collection site (**Figure 6E-F**; **Supplemental Table S20**). Analyses were restricted to patients with complete data for all three measures and the adjustment covariates. Among patients younger than 70, immune age acceleration (HR = 2.24, p = 0.003), LDS-5 (HR = 1.37, p = 0.030), and ECOG (HR = 1.45, p = 0.028) were each associated with shorter OS (n = 201, 56 events; **Figure 6E**). For PFS, immune age acceleration (HR = 1.63, p = 0.017) and ECOG (HR = 1.53, p < 0.001) remained significant, while LDS-5 did not (HR = 0.97, p = 0.809; n = 198, 105 events; **Figure 6E**). In patients greater than age 70, only ECOG was associated with shorter OS (HR = 1.48, p = 0.046; n = 95, 52 events; **Figure 6F**), whereas for PFS, both LDS-5 (HR = 1.38, p = 0.025) and ECOG (HR = 1.44, p = 0.022) were associated with outcome, but immune age acceleration was not (HR = 0.96, p = 0.886; n = 95, 73 events; **Figure 6F**). Thus, in younger patients, immune age acceleration remained associated with both OS and PFS after adjustment for conventional measures of fitness, namely ECOG and LDS-5.

We then compared models containing each measure individually with those including all three measures to determine whether adjustment for other fitness measures altered their associations with outcomes (**Figure 6G**). Associations that were significant in the individual models remained significant after joint adjustment. The association between immune age acceleration and outcome also differed by age group, with significant interactions for both PFS (interaction HR ratio = 1.84, p = 0.018) and OS (interaction HR ratio = 2.58, p = 0.004). In contrast, no significant age-group interactions were observed for ECOG or LDS-5 (**Supplemental Table S19**), indicating the age-dependent prognostic association was specific to immune age acceleration.

Finally, we added ISS stage to the multivariable models to evaluate whether these associations persisted (**Supplemental Figure 6D-6E**). In patients under age 70, immune age acceleration remained associated with shorter OS (HR = 2.21, p = 0.005), whereas LDS-5 (HR = 1.10, p = 0.566) and ECOG (HR = 1.35, p = 0.083) were no longer significant. For PFS, immune age acceleration (HR = 1.52, p = 0.045) and ECOG (HR = 1.45, p = 0.006) remained associated with outcome in the complete-case cohort (n = 198). However, when this PFS analysis was extended to permit patients with missing ECOG or LDS-5 measures (n = 229, 127 events), immune age acceleration was no longer significantly associated with PFS (HR = 1.34, p = 0.102), and a model using missing-indicator terms for ECOG and LDS-5 produced a similar nonsignificant estimate (HR = 1.37, p = 0.075). The 31 excluded patients from the complete-case PFS analysis did not differ from included patients in immune age acceleration (Mann-Whitney U, p = 0.77), and no significant interaction between immune age acceleration and exclusion status was observed (p = 0.589). Collectively, these results illustrate that immune age acceleration captures prognostic information distinct from functional status, laboratory deficits, and disease stage, with persistent association with shorter OS in younger patients, consistent with its lack of correlation with ECOG and weak correlation with LDS-5.

## DISCUSSION

In this study, we developed immune aging clocks for ten immune and progenitor cell types using publicly available single-cell BM and PB data collected from healthy donors, calibrated the clocks to healthy BM samples, and applied them to a single-cell BM MM dataset consisting of 337 patients with NDMM. In patients younger than 70 years old at diagnosis, those who had an immune system that appeared older than expected experienced shorter OS, independent of standard clinical risk factors including tumor burden, performance status, and laboratory deficits. Beyond their prognostic value, the immune aging clocks allowed us to investigate aging-related transcriptional patterns associated with disease progression in younger MM patients. Patients with accelerated immune aging showed stronger inflammatory signaling, a shift towards terminally differentiated T effector states, and changes in interferon and metabolic programs. At first progression, isolated monocyte aging demonstrated a significant risk of decreased continued survival and was correlated with monocyte-driven immune suppression. Together, these findings suggest that immune age captures a significant dimension of clinical and biological heterogeneity in MM that is incompletely captured by chronological age, disease stage, or functional status.

Inflammatory signaling was the most consistent feature of immune aging across cell types. TNF-α/NF-κB signaling increased with predicted age in every cell type in the MM cohort except for HSCs and in all ten cell types in healthy donors, consistent with inflammaging as a core feature of immune aging.^24^ However, a per-patient NF-κB score was not associated with survival, and the clock remained associated with OS after accounting for NF-κB enrichment. Inflammation is therefore a part of what the clock measures but is not by itself sufficient to capture risk in fast agers. Interferon signaling also shifted toward weaker type I and stronger type II responses in fast agers, consistent with the pattern linked to rapid progressors in Pilcher et al.^18^ This inflammatory shift was reinforced at the intercellular level, where memory lymphocytes promoted TNF, CCL3, and IFN-γ signaling cascades in fast agers, leading to increased NF-κB activation and inflammation across recipient compartments. Whether this inflammatory environment is a cause or consequence of MM remains unclear, as we cannot determine whether MM accelerates immune aging, whether an aged immune system allows MM to progress, or an interaction between the two. Immune age acceleration showed a weak relationship with conventional measures of tumor burden, suggesting age acceleration is not a simple consequence of disease burden.

Another clear feature of accelerated immune aging was an increase in differentiated T effector cells. Patients with the most accelerated T cells had greater abundance of effector and cytotoxic T cells and fewer central memory cells. Their memory T cells also expressed greater enrichment of terminally differentiated markers such as *B3GAT1* (CD57), *ZEB2*, and *GZMB* while losing expression of *CD28*, *TCF7*, and *CCR7*, which are needed for co-stimulation and renewal of memory T cells. Similar features of increased exhausted and senescent T cells have been described in MM.^33^ Expansion of cytotoxic CD4^+^ T cells has also been observed in older individuals.^34^ Even with these abundance changes, in CD4^+^ memory T cells, most of the association between age acceleration and survival came from transcriptional changes within subtypes rather than compositional shifts. In CD8^+^ memory T cells, transcriptional changes still contributed to half of the OS association, with the other 50% coming from compositional changes and the interaction between compositional and transcriptional changes within CD8^+^ subtypes. The increased mitochondrial programs in the most accelerated CD4^+^ memory T cells also fit with the higher mitochondrial capacity described in senescent CD4^+^, but not CD8^+^ T cells in Callender et al.^26^

Accelerated NK cells showed a parallel shift from conventional to adaptive NK subtypes. Increased adaptive NK abundance is linked to CMV infection, and CMV is also thought to drive age-related increased abundance of differentiated T effector cells.^29, 35^ Since CMV serology was not available for this cohort, we could not test the link between CMV infection and immune age acceleration, but the NK shift suggests that our immune age clock may partially reflect CMV infection. Other exposures that accelerate immune aging were also not fully characterized. Obesity and type 2 diabetes are associated with accumulation of senescent T cells, smoking has persistent effects on adaptive immune responses, and clonal hematopoiesis of indeterminate potential (CHIP), which is present in approximately one in five patients with MM undergoing ASCT, is associated with shorter survival in patients not receiving immunomodulatory maintenance.^36–39^ Although BMI and glucose were included in LDS-5, which only weakly correlated with immune age acceleration (ρ = 0.17, q = 0.056), diabetes, smoking history, and CHIP status were not evaluated. Future studies incorporating CMV serology, comorbidity status, smoking and CHIP data will help clarify the contributions of these exposures to immune age acceleration.

Although the cell type clocks were highly correlated with one another at diagnosis, they became less coupled after treatment and first progression, and changes in monocyte age acceleration at first progression were associated with subsequent survival. While this outcome association is only based on 17 events from 44 patients, it provides initial evidence that myeloid cells shape MM progression. Myeloid cells were previously found to be enriched in rapidly progressing patients in this cohort,^18^ consistent with the associations between monocyte abundance and overall survival we observed at baseline. At progression, monocytes that aged more than expected showed activation of IL-6-JAK-STAT3, interferon, and NF-κB signaling, together with higher expression of inhibitory receptors, including *LILRB2*, *CD300A*, and *SIGLEC10*. Previous work has demonstrated that *SIGLEC10* recognizes CD24 on tumor cells and can limit macrophage phagocytosis of tumor cells, raising the possibility that aged monocytes are less able to clear MM cells.^40^ Collectively, our findings suggest that monocyte aging at first progression may contribute to an immunosuppressive bone marrow environment, and the cell type clocks can capture changes after treatment that a single global score misses.

Immune age acceleration was only associated with outcomes in patients younger than age 70. The interaction between immune age acceleration and age group was significant, indicating a true difference between age groups, although several possible explanations for this difference in age groups remain. First, the clocks may be less precise in older patients. Fewer healthy donors older than age 80 were available for training, and because many patients already showed positive acceleration, older patients whose immune cells appear much older than their chronological age may fall outside the range that the clocks were trained to capture. Second, the relationship between immune age acceleration and survival may be nonlinear, with diminishing prognostic differences as immune age increases. Third, in older patients, factors other than immune age may dominate outcomes. In patients over age 70, chronological age, ECOG performance status, and laboratory deficits were associated with survival, and these patients received less ASCT and triplet therapy. Consequently, comorbidities, frailty, and treatment intensity may outweigh the contribution of immune aging such that more accelerated aging minimally influences outcomes past a certain threshold. Further validation in larger, age-diverse cohorts will be necessary to distinguish these potential explanations and determine whether the prognostic relevance of immune age acceleration varies continuously with chronological age.

Immune age acceleration was also more consistently associated with poorer OS than PFS in patients younger than age 70, suggesting that immune fitness may have greater relevance to outcomes over the course of disease rather than the timing of first progression. Progression is driven largely by tumor biology and response to first-line therapy, whereas OS can additionally reflect the ability to control relapsed disease, withstand repeated treatment, and combat infections, a major cause of death in MM,^41^ all of which may depend on immune fitness. This is consistent with our monocyte finding, in which greater monocyte age acceleration was also associated with shorter OS after first progression. Immune fitness also has implications for treatment planning. CAR-T cell products enriched for naïve and less differentiated T cells, particularly when collected earlier in the course of disease, have been associated with better expansion and response in MM.^42^ Patients with accelerated immune aging, and whose memory T cells have lost CD28, TCF7, and CCR7, may therefore benefit from earlier T cell collection. Importantly, immune age was also independent of ECOG performance status and quality of life measures and remained associated with OS after adjustment for ECOG and laboratory deficits. As MM is increasingly diagnosed in younger patients,^2^ biological measures of immune fitness may help identify higher-risk patients not captured by current risk assessments.

Our approach offers several methodological strengths. We developed cell type-specific aging clocks using elastic net models rather than deep learning models. Because single-cell data is sparse and can vary between batches, more complex models can easily learn technical noise rather than biology. Moreover, elastic net models offer greater interpretability, allowing us to examine which genes each clock relies on and revealing that these genes reflect established features of immune aging. Our models also allow us to reduce the clocks to smaller sets of marker genes, providing a foundation for developing targeted assays for clinical assessment, such as flow cytometry, to identify patients with aged immune systems. However, because elastic net selects among correlated genes, individual coefficients reflect what the model uses rather than what causes aging, so markers of age acceleration should be examined across datasets before deciding on potential markers.

This study also has limitations that warrant consideration. Healthy bone marrow scRNA-seq data is limited, and the clocks were trained on 86 bone marrow samples supplemented with peripheral blood. Additional healthy bone marrow samples, especially from older donors, would improve accuracy and help establish how much immune aging varies among healthy individuals. Because the reference atlas was annotated into ten broad cell populations, clocks were trained at the lineage level, and our subsequent subtype-specific analyses were exploratory. Furthermore, the MM cohort was profiled using 3′ scRNA-seq without TCR or BCR sequencing, preventing direct assessment of clonal expansion, a central feature of T cell aging. Finally, because the clocks were trained on both blood and bone marrow, future studies with paired blood and bone marrow samples could test whether a blood-based version captures the same risk, which would allow immune age to be measured from a blood draw. Future studies incorporating functional immune assessments, larger longitudinal cohorts, and paired BM and peripheral blood samples will be important for validating these findings and translating them into a prognostic immune aging assay.

In summary, cell type-specific immune aging clocks identify younger patients with MM whose immune cells exhibited greater age acceleration and who experienced shorter survival, independent of chronological age, disease stage, and functional status. Beyond prognosis, these clocks revealed inflammatory signaling and effector differentiation of T cells within the BMME, as well as outcome-associated changes in monocytes at progression. Together, these findings establish immune age acceleration as a potential biomarker of clinical and biological heterogeneity in MM and support further investigation of its relevance to risk stratification and treatment planning, particularly as T cell-directed therapies are increasingly incorporated into earlier lines of treatment.

## Notes

### Competing Interest Statement

The authors have declared no competing interest.

